# PEA polymer-coated nanotopography delivers solid-state BMP2, enhances mesenchymal stem cell adhesion, prevents bacterial biofilm formation and protects cells from quorum sensing virulence factors

**DOI:** 10.1101/2020.09.17.302455

**Authors:** Laila A. Damiati, Monica P. Tsimbouri, Mark Ginty, Virginia Llopis Hernandez, Peter Childs, Vineetha Jayawarna, Yinbo Xiao, Karl Burgess, Julia Wells, Mark R. Sprott, R.M. Dominic Meek, Peifeng Li, Richard O.C. Oreffo, Angela Nobbs, Gordon Ramage, Bo Su, Manuel Salmeron-Sanchez, Matthew J. Dalby

**Affiliations:** Centre for the Cellular Microenvironment, Institute of Molecular, Cell and Systems Biology, College of Medical, Veterinary and Life Sciences, University of Glasgow, Glasgow G12 8QQ, UK; Department of Biology, Faculty of Biological Science, University of Jeddah, Jeddah 23890, SA; School of Oral and Dental Sciences, University of Bristol, Bristol, BS1 2LY, UK; Centre for the Cellular Microenvironment, Division of Biomedical Engineering, School of Engineering, University of Glasgow, Glasgow G12 8LT, UK; Department of Biomedical Engineering, University of Strathclyde, Glasgow G1 1QE, UK; Glasgow Polyomics, College of Medical, Veterinary and Life Sciences, University of Glasgow, Switchback Rd, Bearsden, Glasgow G61 1BD, UK; Bone and Joint Research Group, Centre for Human Development, Stem Cells and Regeneration, Institute of Developmental Sciences, University of Southampton, Southampton SO16 6YD, UK; Department of Orthopedics, Queen Elizabeth II University Hospital, Glasgow G51 4TF, UK; Oral Sciences Research Group, Glasgow Dental School, School of Medicine, College of Medical, Veterinary and Life Sciences, University of Glasgow, Glasgow, G12 8TA, UK

**Keywords:** Multifunctional materials, nanotopography, nanoscale coatings, mesenchymal stem cells, bactericidal, *Pseudomonas aeruginosa*, quorum sensing and signalling molecules

## Abstract

Post-operative infection is a major complication in patients recovering from orthopaedic surgery. As such, there is a clinical need to develop biomaterials for use in regenerative surgery that can promote mesenchymal stem cell (MSC) osteospecific differentiation and that can prevent infection caused by biofilm-forming pathogens. Nanotopographical approaches to pathogen control are being identified, including in orthopaedic materials such as titanium and its alloys. These topographies use high aspect ratio nanospikes or nanowires to prevent bacterial adhesion but these features puncture adhering cells, thus also reducing MSC adhesion. Here, we use a poly(ethyl acrylate) (PEA) polymer coating on titanium nanowires to spontaneously organise fibronectin (FN) and to deliver bone morphogenetic protein 2 (BMP2) to enhance MSC adhesion and osteospecific signalling. This nanotopography when combined with the PEA coating enhanced osteogenesis and reduced adhesion of *Pseudomonas aeruginosa* in culture. Using a novel MSC–*Pseudomonas aeruginosa* co-culture, we also show that the coated nanotopographies protect MSCs from cytotoxic quorum sensing and signalling molecules. We conclude that the PEA polymer-coated nanotopography can both support MSCs and prevent pathogens from adhering to a biomaterial surface, thus protecting from biofilm formation and bacterial infection and supporting osteogenic repair.

---

For regenerative medicine to progress, to support an ageing population and to improve the treatment of traumatic injuries, we need to develop multifunctional biomaterials that can better guide and support tissue regeneration. Most biomaterials are designed to initiate a specific regenerative response, for example, in mesenchymal stromal or stem cells (MSCs), which support the differentiation of osteoblasts into bone.^1^ However, as well as considering the bony fixation of an implant, clinicians also need to control infection. Surgical site infection for orthopaedic procedures in the UK is up to 10%, depending on reporting, although these are mainly superficial rather than deep infections.^2^ For procedures such as neck of femur fracture arthroplasty, which is typically performed on elderly and frail patients, mortality within 12 months is at 10-40% of surgical patients, with post-operative infection being a major complication.^3^ Indeed, aseptic loosening from poor bone integration and infection are the two leading causes of orthopaedic implant failure, accounting for 18% and 20% of re-surgeries for total knee arthroplasty, respectively.^4^

Thus, materials that can perform duel functions, such as concurrently enhancing bone healing and reducing infection, would help to improve the outcomes of orthopaedic surgical patients. Indeed, such materials are of particular importance in light of the increased risk of infection as bacteria gain resistance to antibiotics, which could potentially render surgery less routine.^5^

Bone regeneration materials have been researched since the osteoconductive potential of hydroxyapatite and bioglasses was first reported in the 1970’s.^6, 7^ Since then, the osteoinductive potential of materials has been modulated by altering substrate stiffness,^8, 9^ as well as their viscoelasticity,^10, 11^ chemistry^12^ and topography^13^, as assessed through changes in MSC mechanotransduction.^14–18^ Osteoinductive soluble factors, such as bone morphogenetic protein 2 (BMP2), have also been investigated using biomaterial platforms. The delivery of such potently regenerative factors is important as they have short half-lives and systemic, as well as targeted, effects.^19^ For example, one delivery system called InFuse employs collagen sponges to deliver BMP2 to intervertebral discs being fused. In this system, a supraphysiological dose of BMP2 is delivered in a burst-like manner to provide a high concentration at the target site. However, while this BMP2 dose supports the efficient fusing of bone, it produces off target effects, such as ectopic bone formation, nerve damage, sterility and even cancer.^20, 21^ As a result, solid-phase, materials-based approaches are being considered as an alternative means to deliver osteoinductive factors more safely.^22^

The solid-phase delivery of growth factors, such as BMP2, is considered an important goal because when *in vivo*, growth factors bind to the extracellular matrix (ECM). The binding of growth factors to the ECM increases their stability, enables low levels of growth factor to effect local regeneration^2^, and increases their functional lifespan.^23–25^ Approaches that aim to mimic the binding of growth factors to the ECM include: the fabrication of positively and negatively charged layers with growth factors trapped in between each layer;^26, 27^ the inclusion of ECM components that have affinity for growth factors, such as heparin sulphate proteoglycans, fibronectin (FN) or fibrinogen;^28^ and the direct binding of growth factors on to materials.^24, 29–31^

Solid-phase growth factor delivery can be further enhanced through synergy with integrins. Integrins are transmembrane receptors that bind cell-adhesion motifs, such as the arginine-glycine-aspartic acid (RGD) motif, which are found in ECM proteins such as FN.^32–35^ When integrins bind to ECM ligands, it drives the formation of focal adhesions, which can effect osteogenic signalling by sequestering focal adhesion kinase (FAK). FAK drives cytoskeletal contraction through Rho A kinase (ROCK) and stimulates extracellular signal related kinase (ERK 1/2), leading to the activation (via phosphorylation) of the osteogenic master transcription factor, runt-related transcription factor 2 (RUNX2).^14, 15, 36–39^ BMP2 can, non-canonically, act through ERK 1/2 to effect osteogenesis, and can also canonically activate RUNX2-mediated osteogenesis through SMAD (small mothers against decapentaplegic) signalling.^19^ Indeed, ECM molecules, such as FN and laminin, contain integrin-binding motifs and growth factor-binding regions in close apposition, and this can drive enhanced regeneration.^40, 41^ FN itself is a dimeric glycoprotein that has two subunits linked by a single disulphide bond; these two subunits contain three repeating modules (types I, II and III), which mediate FN’s interaction with other FN molecules. These interactions facilitate the formation of protein networks consisting of FNI_1-5_ and FNIII_1-2_, integrins (FNIII_9-10_ containing RGD) and growth factors (FNIII_12-14_ forming a heparin binding domain).^42^ The structure of FN appears designed to provide signalling synergy that, as bioengineers, we could exploit.

Using antibiotics loaded into materials as a means of treating deep infections poses problems due to poor release kinetics and lowered mechanical properties of the biomaterials used to deliver them. For example, antibiotics can be loaded into cement spacers but these spacers often require exchanging and are mechanically weak.^43^ Other approaches, such as adding a silver coating to a material, can reduce bacterial attachment but the high doses required to kill pathogens also reduce osteoblast activity.^44^ Topographical approaches offer another interesting way to address this problem. One approach has been to mimic the bactericidal, high aspect ratio topographies that are present on insect wings, such as on Clanger cicada (*Psaltoda claripennis*)^45^ and the dragonfly (*Diplacodes bipunctata*)^46^. Materials such as black silicon^46^ and titanium (a widely used orthopaedic material)^47^ have been used to mimic this affect by shaping them into topographies that produce high levels of mechanical deformation of pathogen membranes, leading to bacterial cell rupture and death.^45, 46, 48^ However, these very spiky topographies are far from optimal in terms of osteogenesis.^47^ Use of integrin mimics has been shown to enhance bioactivity,^49^ but does not provide robust osseoinduction potentially because only one facet of the extracellular matrix, adhesion, is being considered.

Here, we investigate whether adding integrin and BMP2 functionality to a high aspect ratio topography formed in titania, the oxide coating of titanium and its alloys, will provide both an anti-bacterial and osteogenic effect for use in orthopaedic materials. To achieve this, we added a thin (<60 nm) coating of the polymer poly(ethyl acrylate) (PEA) to the topography; this coating thickness is important because the sharpness of features is believed to be central to the implants antibacterial activity.^45, 46, 48^. PEA has been shown to cause FN to open up to reveal its integrin binding site (RGD at FNIII9-10) and its heparin binding region, which can be decorated with BMP2 and which drives MSC osteogenesis in synergy with integrin binding.^19^ This opening of FN is important as typically FN absorbs to biomaterials in a globular manner with its binding sites inaccessible to cells until they have opened FN out.^19, 40^

To investigate the osteogenic and antibacterial properties of a PEA-coated titania topography, we used MSCs and the bacterial pathogen *Pseudomonas aeruginosa* (*P. aeruginosa*), which causes sepsis associated with orthopaedic infection.^50^ *P. aeruginosa* was also selected because it is a Gram-negative bacteria and thus presents a good first target due to its cell wall structure. First, we identified nanotopographies with a PEA nanocoating that could deliver low doses of BMP2 able to both enhance MSC osteogenesis and to reduce biofilm formation in culture. We then used a novel MSC and *P. aeruginosa* co-culture to show that the PEA-coated nanotopographies can greatly reduce bacterial colonisation and MSC susceptibility to cytotoxic quorum sensing molecules. Finally, we generated 3D porous Ti scaffolds coated with PEA and with a Young’s modulus similar to bone to show translation into 3D.

## Results and Discussion

### Coating Ti nanowires with plasma PEA

In this study, we used nanospike topographies made in the oxide layer of Ti. To generate the nanospike topographies, we used Ti discs that were polished to a mirror finish and washed using sonication. We then generated high aspect ratio nanowires on the Ti discs using an alkaline hydrothermal process, in which samples were incubated with 1 M NaOH at 240°C for 1 or 2 hrs; the longer the incubation time, the longer the wires that grow. SEM and AFM were used to show topography formation with average heights of 400 nm and 550 nm for the 1 and 2 hr samples, respectively (Figure 1 a, b). Surface roughness was measured by AFM, showing a stepwise increase in Ra value, from the flat control to the 1- and 2-hr samples, as expected (Figure 1c).

**Figure 1.**
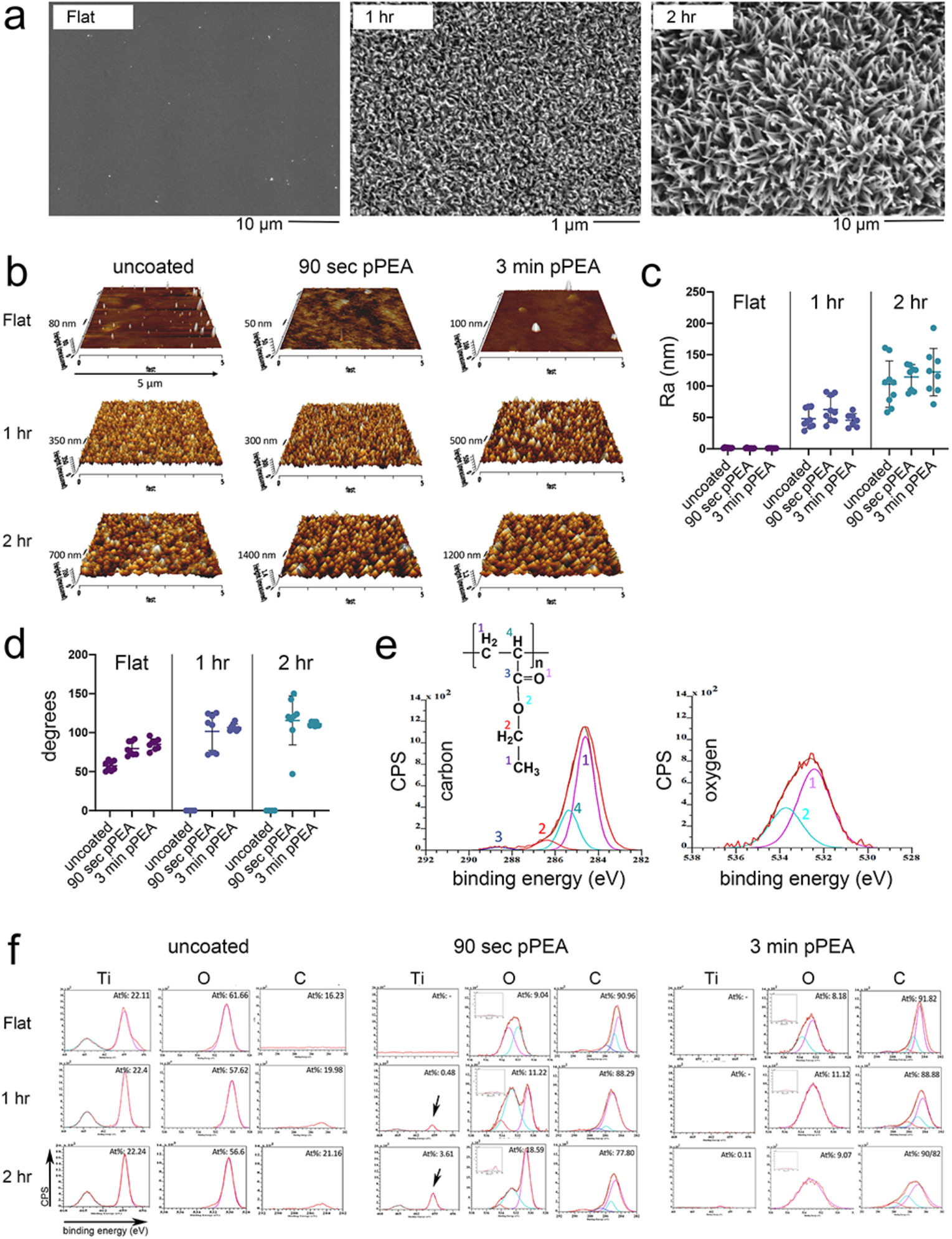
Formation of nanospike topographies and PEA coating of surfaces. (a) SEM images of the flat control, 1-hr nanospikes and 2-hr nanospikes, showing the formation of high aspect ratio nanofeatures of increasing size. (b) AFM images of the uncoated controls (flat, 1-hr and 2-hr topographies), and of the 90-sec and 3-min pPEA-coated topographies (flat, 1-hr and 2-hr). The results show that the topographies remain evident after the PEA-coating process. (c) AFM Ra measurements, demonstrating that an increasing topography size increases Ra, and that the PEA coating does not change a topography’s Ra values (mean±SD, n=8). (d) Sessile drop contact angle measurements, showing increased hydrophobicity with the addition of the PEA coating (mean±SD, n=8). (e) XPS spectra generated from PEA reference (binding compositions are peak fitted onto the image). (f) XPS measurements of uncoated Ti controls with and without topography, showing the presence of Ti and O. XPS spectra for the 90-sec and 3-min PEA coatings show that with the shorter, 90-sec coating, both Ti- and PEA-related peaks are present. With the longer, 3-min PEA coating, only PEA-related peaks were observed. The data show that high aspect ratio nanotopographies were successfully fabricated and coated with pPEA.

Plasma polymerisation of the ethyacrylate (EA) monomer was used to create PEA coatings on the nanotopography surfaces. SEM and AFM imaging, and RA value calculations, showed that features retained their original, pre-coated topographies after being coated with PEA for 90 seconds or 3 minutes at 100 W power (Figure 1 b, c; Supplementary Figure 1). PEA is a hydrophobic polymer, and so contact angle measurements of the topographies were taken and showed an increase in hydrophobicity with coating; it was not possible to take contact angle measurements of uncoated topographies (Figure 1d). X-ray photoelectron spectroscopy (XPS) was used to show the successful polymer coating of the topographies. Figure 1e shows typical carbon and oxygen traces for PEA and Figure 1f shows spectra for uncoated and coated samples. Without coating, Ti and O peaks were visible for all samples, reflecting the topography being formed in the passivating TiO_2_ layer. With the shorter, 90-sec PEA coating, peaks relating to PEA were noted on all samples but no Ti peak could be observed on the flat control samples, while small Ti peaks were evident on the 1- and 2-hr topographies, especially for the larger features on the 2-hr sample. In samples coated with PEA for the longer 3-min period, the Ti peaks disappeared on all samples while peaks relating to PEA were observed (Figure 1f) (note that larger XPS images are provided in Supplementary Figure 2). This data indicates that with 90-sec PEA coating a non-conformal PEA layers are formed, especially with the larger features, with areas of Ti still apparent. However, with the with 3-min coating, the PEA layer has completely covered the Ti surface on all samples.

### Identifying an optimal osteogenic nanotopography and coating combination

We next screened MSCs plated on all the test surfaces for 21 days, with and without FN/BMP2, using rapid qPCR to screen for the following osteoblast-related transcripts: osteocalcin (OCN), osteopontin (OPN) and osteonectin (OCN). This was done to select a reasonable number of surfaces for further in-depth characterisation and for bacterial screening. For the solid-phase presentation of BMP2 on PEA-coated samples, surfaces were loaded with 100 ng/ml BMP2 before culture. To deliver soluble BMP2 (when using uncoated samples), the culture media was supplemented with 25 ng/ml BMP2, which was refreshed every 3 days (175 ng/ml for 21 day culture, 250 ng/ml for 28 day culture and 300 ng/ml for 35 day culture). From these experiments, we selected the 90-sec PEA-coated, 2hr nanowire topography loaded with FN and BMP2 (90 sec pPEA-2h FN/BMP2) to take forward because of the high OCN and OPN values observed compared to other coated nanotopographies (Figure 2a). This topography was next benchmarked to control samples (uncoated-flat, uncoated-flat with soluble FN and BMP2, 90 sec pPEA-flat with FN and BMP2, and 3 min pPEA-flat with FN and BMP2), by qPCR after 28 days of culture (Figure 2b). In this analysis, the selected surface showed higher levels of osteogenic transcript expression relative to the uncoated control sample, even when BMP2 was added to this control, and benchmarked well to flat control pPEA coated samples with solid phase BMP2 presentation via FN.

**Figure 2.**
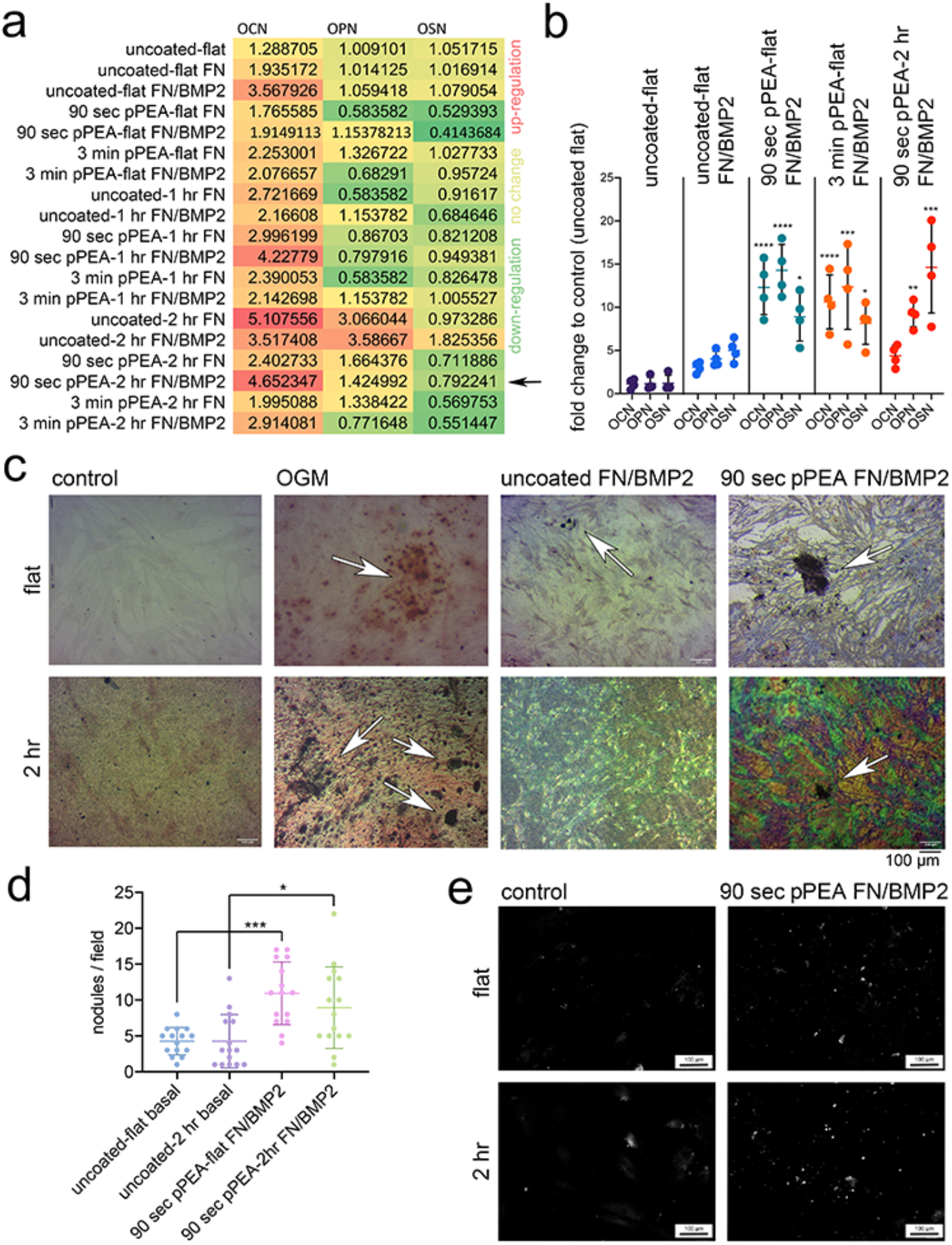
Analysis of MSC osteogenesis on nanowire topographies. (a) Osteocalcin (OCN), osteopontin (OPN) and osteonectin (OSN) transcript levels, shown as a heatmap and assayed by rapid qPCR screening of MSCs after 21 days of culture on different nanowire topographies (as shown). These results highlighted the 2 hr nanowires with 90 sec pPEA coating (indicated by a black arrow) as being optimal for osteogenesis on high aspect ratio nanofeatures (results show mean values, d=1, r=4, t=2 (n= number of MSC donors, r = material replicates, t = technical replicates), red=up-regulation, yellow=no change, and green = down-regulation). (b) OCN, OPN and OSN transcript levels in MSCs cultured for 28 days, as assayed by qPCR. MSCs cultured on flat surfaces show similar levels of MSC osteogenesis in the presence or absence of soluble FN/BMP2. MSCs cultured on PEA-coated, 2 hr nanowires in the presence of soluble or solid-phase BMP2 show enhanced osteogenesis relative to uncoated flat controls (graph shows mean±SD, d=1, r=4, t=2, stats by ANOVA and Tukey test where *=p<0.05, **=p<0.01, ***=p<0.001 and ****=p<0.0001). (c) Alizarin red staining (showing mineralisation) in MSCs cultured for 35 days on the indicated topographies shows that 90 sec PEA coating with FN and BMP2 has a similar mineralisation profile to use of osteogenic media on planar titanium surfaces while use of soluble FN and BMP2 confers no advantage to planar Ti. Arrows indicate mineralized nodule formation. (d) Quantification of calcein blue staining for the number of bone nodules formed in MSCs after 35 days in culture on the indicated topographies (graph shows mean±SD, d=1, r=4, t=15, stats by ANOVA and Tukey test where *=p<0.05 and ***=p<0.001). (e) Representative images of calcein blue fluorescent histology, showing less background staining than that produced by Alizarin red. These results indicate that a PEA coating, combined with FN and BMP2, rescues MSC osteogenesis on high aspect ratio nanowires, more effectively than soluble BMP2.

A histological analysis of MSC mineralisation was performed after 35 days of culture using Alizarin red staining. While it was more challenging to see mineralized nodules on the 2-hr topographies, no nodules were observed when MSCs were cultured on the flat or 2-hr Ti surfaces. By contrast, widespread mineralisation was seen in the presence of an osteogenic media (used as a control), while MSCs cultured on the 90 sec pPEA-2h FN/BMP2 surface showed increased mineralisation compared to flat controls, with the PEA coating enhancing the osteogenic capacity of BMP2 compared to use of soluable BMP2(Figure 2c). A comparison of the number of bone nodules produced on the uncoated and 90-sec pPEA surfaces after 35 days of culture using calcein blue staining confirmed the Alizarin red results: that more nodules are seen when the flat control surface or the 2-hr nanospikes were coated with 90 sec pPEA FN/BMP2 (Figure 2d). (Calcein blue was used for quantification as Alizarin red produced too high a background to give reliable quantification, while calcein blue, which is typically used in *in vivo* bone formation assays,^51^ gave background-free fluorescence images (Figure 2e).

Overall, the data show that high aspect ratio nanowires reduce osteogenesis as expected.^47^ However, the use of pPEA coating with FN / BMP2 can rescue osteogenesis, as assessed by osteogenic gene expression and the number of bone nodules formed. It is noteworthy that solid-phase BMP2 delivery (via the PEA-coated nanowires) provided enhanced osteogenesis compared to the soluble delivery of BMP2 (via uncoated nanowires), as assessed by osteogenic gene expression (Figure 2b) and mineralisation assays (Figure 2c), despite significantly more BMP2 being used for soluble delivery.

### Assessing FN/BMP2 interactions with PEA-coated nanowires

As discussed, PEA drives the formation of FN nanonetworks.^19^ These networks are clearly visible when FN is absorbed onto spin-coated PEA, as shown by AFM in Figure 3a, where PEA has been spin coated onto glass and coated with FN. The use of plasma coating causes tighter networks to form, as shown in Figure 3b with some FN morphological differences observed in dependence of plasma energy (J (joules) = W (Watts) x s (time)). For the same amount of FN absorbed onto the surface, the PEA-driven network formation drives enhanced availability of the FN integrin-binding site, RGD (HFN7.1), and of the heparin-binding domain (which binds to many growth factors) and also increases the amount of BMP2 absorbed onto the surface, compared to FN and BMP2 added to glass coverslips without PEA coating (Figure 3b).

**Figure 3.**
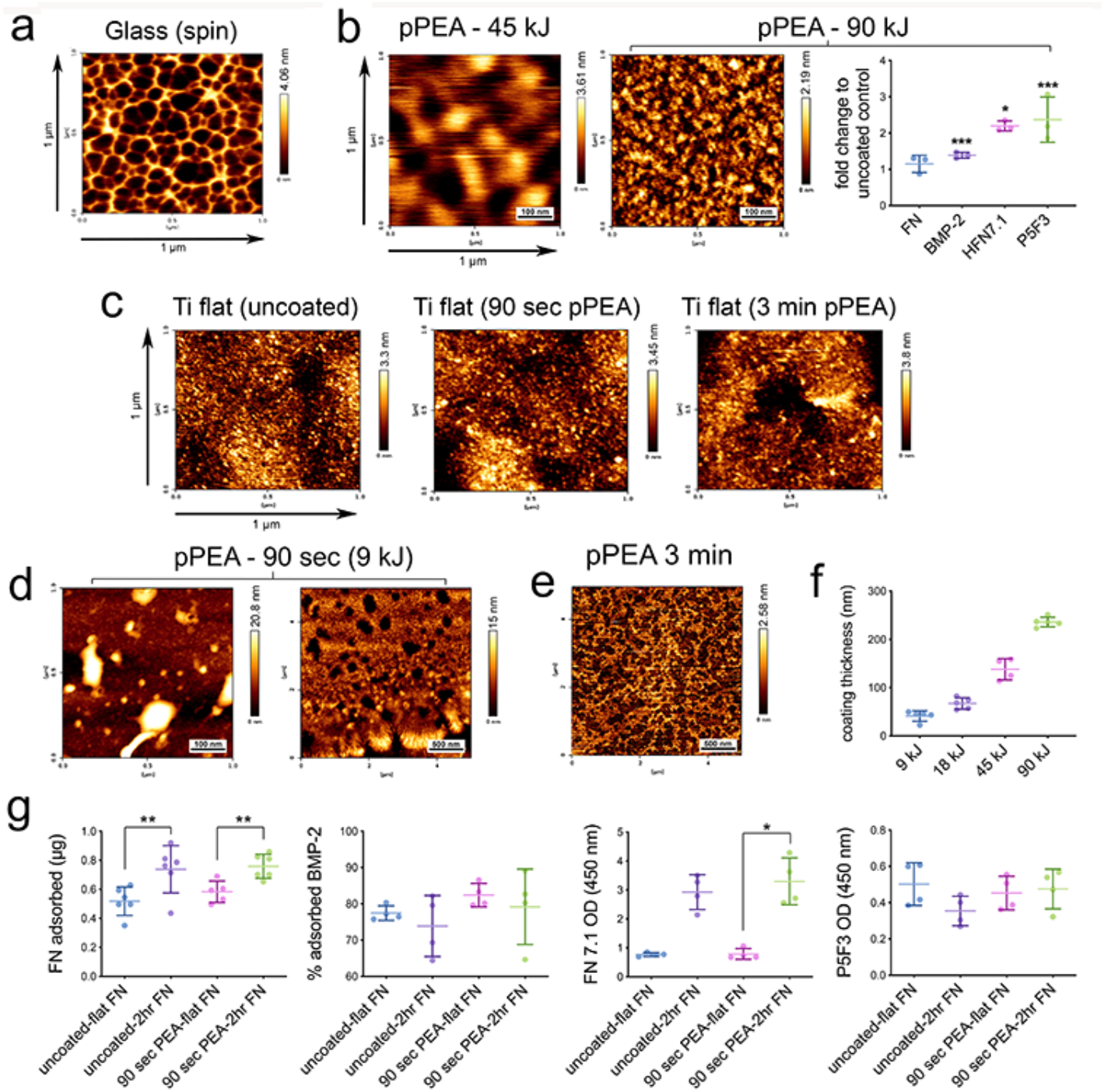
Evaluation of FN and BMP2 interactions with pPEA. FN nanonetworks were assessed by AFM. (a) FN nanonetwork on a glass coverslip with spin-coated PEA, showing a diffuse network. (b) FN nanonetwork on a glass coverslip with pPEA spin coated at 45 kJ and 90 kJ, showing dense FN networks. (Right) The availability of the RGD (HFN7.1) and heparin binding sites (P5F3) on FN, as assessed by ELISA. These sites’ availability increased with a concomitant increase in BMP2 absorption, when FN was present as a network on 90 kJ pPEA (graph shows mean±SD, n=3, stats by ANOVA and Tukey test where *=p<0.05 and ***=p<0.001). (c) FN networks were hard to image on the nanorough pPEA-coated Ti surfaces. (d) Glass coverslips were used to observe FN interactions. On 90 sec (9 kJ) pPEA coated coverslips, FN formed both globular and fibrillar networks but only fibrillar networks (e) on 3-min (18 kJ) pPEA-coated coverslips. (f) Increasing PEA plasma energy increased the coating thickness (graph shows mean±SD). (g) ELISA was used to observe FN and BMP2 interactions with flat and 2 hr nanowire Ti surfaces, with and without 90 sec pPEA FN/BMP2 coating. From flat to 2 hrs (with or without coating), an increase in FN absorption was seen. From flat to 2 hrs with 90 sec pPEA coating, an increase in RGD availability was seen. No difference in heparin binding site availability or BMP2 absorption was noted. (Graphs show mean±SD, n=3, stats by ANOVA and Tukey test where *=p<0.05 and **=p<0.01)

Observing networks on Ti was challenging. While the mirror-polished Ti surfaces have an Ra of just 1.37±0.59 nm, this surface is rougher and shows greater variance than glass coverslips (0.67±0.03nm).^52^ Thus, FN networks cannot be observed on Ti surfaces (Figure 3c); as such, we used glass as a model surface for the 90-sec and 3-min pPEA coatings to observe FN interactions. The 90-sec coating resulted in areas of both globular and fibrillar FN deposition (Figure 3d). This is likely because the coating is very thin at this energy (9 kJ) at 41.1±10.8 nm (Figure 3f) and might not conformally coat the surface, supporting XPS data showing that Ti can be detected on the 90-sec PEA coated surface (Figure 1f). The 3-min PEA coating (18 kJ, 67.2±11.8 nm) supported fibrillar FN over the whole surface (Figure 3e), which supports the XPS data showing that Ti cannot be detected on the 3-min PEA coated surface (Figure 1f). These very low energies, much lower than those previously investigated for plasma PEA coatings,^53, 54^ were chosen to ensure that the topographical features retain their high aspect ratio. It is also interesting that the heterogeneous globular / fibrillar FN on the 90 sec pPEA coating best supports osteogenesis when applied to the 2 hr nanowires.

Our assessment of FN absorption, BMP2 absorption, and of the exposure of the RGD (FN7.1) and the heparin binding (P5F3) regions, indicate that the 90-sec PEA coating leads to more FN being absorbed with greater availability of the RGD cell adhesion motif (Figure 3g). However, no change in heparin binding domain measurement or BMP2 surface absorbance was seen (Figure 3g).

### Nanowires and pPEA coating reduce bacterial adhesion and metabolism

*P. aeruginosa* was seeded onto the surface of flat control and 2-hr nanowires with and without 90 sec pPEA FN/BMP2 coatings. Bacteria attached to both flat and 2-hr surfaces, although flagella were not observed on the pPEA-coated or uncoated nanowires but were observed on the flat surfaces (Figure 4a). Flagella are used by cells to locomote, and their loss is usually associated with cell stress, such as nutrient deprivation.^55^

**Figure 4.**
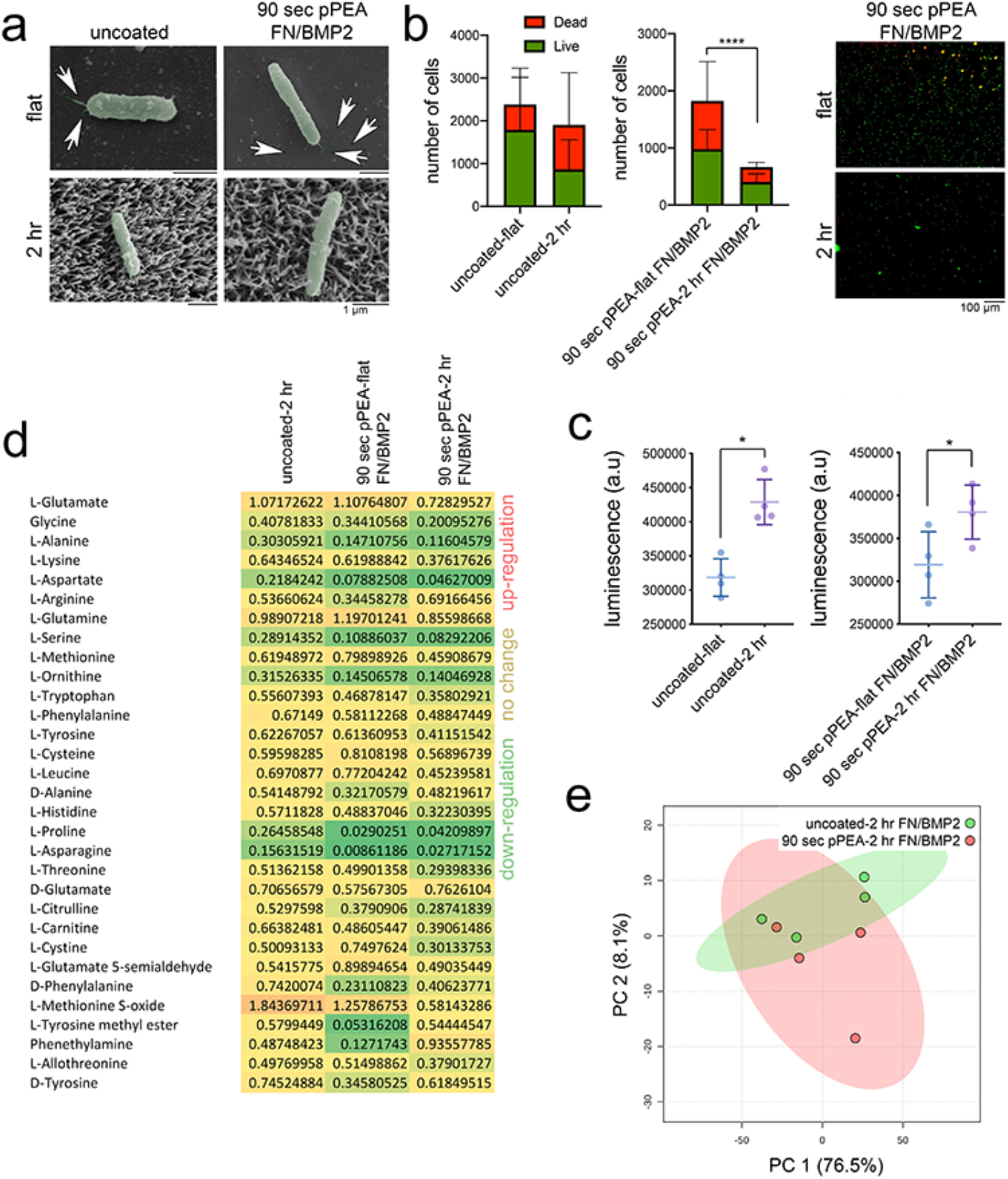
*P. aeruginosa* response to nanowire topography and pPEA coating. (a) SEM images showing that *P. aeruginosa* stays on top of the nanowires and loses flagella (arrows) after 4 hours in culture. (b) Nanowires reduce *P. aeruginosa viability after 8 hours of culture*, especially when the surface is coated with pPEA (graphs show mean±SD, r=3, t=12, stats by T-test where ****=p<0.0001); representative live/dead images shows (right) (grre=live/red=dead). (c) ATP assay for planktonic bacteria, showing that there are more unattached bacteria after 24 hours of culture on 90-sec nanowires relative to flat equivalents (graphs show mean± SD, r=4, t=3, stats by T-test where *=p<0.05). (d) Untargeted metabolomic analysis of amino acids after 24 hours of bacterial culture, showing that nanowires and PEA coating reduced the abundance of amino acids within the attached, sessile bacteria (results show mean values, r=4, t=1, red=up-regulation, yellow = no change, and green = down-regulation). (e) PCA showing that amino acid metabolism is different in *P. aeruginosa* cultured on the 2 hr nanowires, with and without pPEA coating (results show mean values, r=4, t=1). The data shows that both the topography and the pPEA coating reduce bacterial attachment and metabolism.

*P. aeruginosa* viability was also slightly reduced when the bacteria were cultured on uncoated nanowires, relative to uncoated flat controls (Figure 4b). When cultured on pPEA-coated nanowires, *P. aeruginosa’s* viability was significantly reduced, as compared to when the bacteria were cultured on pPEA-coated flat Ti surfaces. This reduction in cell viability was confirmed by fluorescence microscopy (Figure 4b).

We next used an ATP assay to quantify planktonic cells (cells in suspension) to assess *P. aeruginosa*’s attachment to the different topographies. The cultures were inoculated with similar titres of bacteria and so an increase in planktonic cells indicates a reduction in attached, sessile cells that can form biofilms. Our results show that ATP measurements increased in nanowire cultures relative to the flat-surface cultures, for both uncoated and pPEA coated samples (Figure 4c. This indicates that *P. aeruginosa* attachment is reduced on the nanotopographies.

Using untargeted metabolomics^56^ to study *P. aeruginosa* amino acid metabolism in response to the test surfaces, a reduction in identified metabolites was observed for bacteria cultured on the uncoated 2 hr nanowire surface (Figure 4d, relative to uncoated flat controls. This reduction increased with the use of the pPEA FN/BMP2 coating on the flat Ti samples, and became more pronounced on the pPEA FN/BMP2 coated nanowire samples (Figure 4d). This reduction in metabolites represents a likely drop in bacterial activity on the pPEA and nanowire coatings. It is noteworthy that the addition of soluble FN/BMP2 to the bacterial cultures increased the production of some amino acids and so could cause a potential increase in bacterial activity (Supplementary Figure 3). A principle component analysis (PCA) of amino acid metabolism revealed distinct patterns of metabolism in uncoated versus pPEA-coated nanowires, highlighting that surface topography and the PEA coating provide different contributions to the bacterial response (Figure 4e).

### MSCs and nanotopography synergistically decrease bacterial adhesion

In order to create an MSC and *P. aeruginosa* co-culture, we first had to control the very rapid growth of *P. aeruginosa* to allow us to observe the slower MSC response. To achieve this, we used short culture times. MSCs were seeded and cultured for 24 hours to allow their attachment and were then inoculated with *P. aeruginosa* (at 10^3^ CFU/ml) and cultured for 24 hours. The culture media was also supplemented with low doses (0.3%) of penicillin and streptomycin to slow bacterial growth without killing the bacteria (Supplementary Figure 4). After 24 hours of co-culture on uncoated and pPEA-FN/BMP2-coated flat controls, biofilm formation around the attached MSCs was observed by SEM (Figure 5a) and by fluorescence microscopy (Figure 5b). However, when MSCs were cultured on the uncoated and pPEA-FN/BMP2-coated 2-hr nanowires, they spread on the material surfaces and very few attached bacteria were observed (Figure 5 a, b). When *P. aeruginosa* was co-cultured on the nanowires with MSCs, the morphology of their cells appeared deflated and punctured, as previously described for other bacteria cultured on high aspect ratio topographies (Figure 5c).^45, 46, 48^ After 24 hours of co-culture, immunofluorescent staining for phosphorylated, active RUNX2 (pRUNX2), actin and DAPI was used to quantify the attachment and osteogenic differentiation of MSCs cultured on the pPEA-coated flat Ti surface (on which *P. aeruginosa* formed biofilms) and on the 2 hr-nanowire topographies (where bacterial attachment was minimal). Imaging (Figure 5d) and image analysis (Figure 5e, f) showed that more MSCs attached to the coated 2hr nanotopographies relative to flat equivalents and that MSCs on the topographical surfaces expressed higher levels of pRUNX2 relative to flat equivalents. RUNX2 was selected as a marker as it is expressed very early on in MSC osteocommittment.^57^

**Figure 5.**
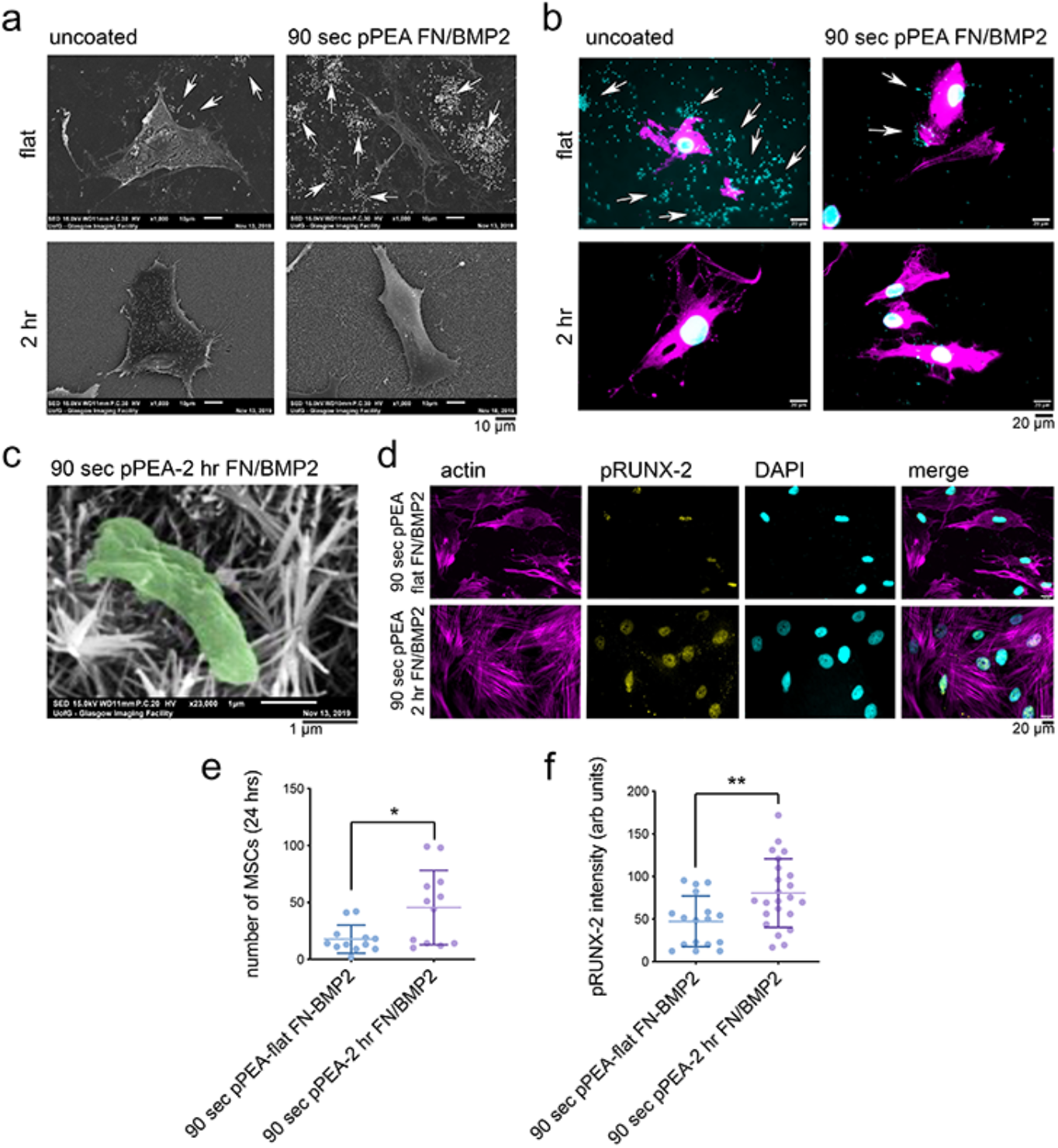
*P. aeruginosa* and MSC co-culture response to nanowire topography and pPEA coating. (a, b) While biofilms could form around MSCs on both uncoated and pPEA-coated flat control surfaces, very few bacteria were seen with MSCs on uncoated and pPEA-coated nanowires (in the fluorescent images, bacteria = cyan, MSCs = magenta). (c) *P. aeruginosa* cultured on the 2 hr nanowires have a deflated, punctured morphology. (d) pRUNX2/actin/DAPI staining showed that MSCs cultured on the 2 hr nanowire samples spread more extensively than those cultured on the indicated flat surface and expressed higher levels of nuclear pRUNX2. (e) Image analysis of MSC number showed increased MSC attachment/growth on the nanowire surfaces. (f) Image analysis showed that MSCs cultured on the nanowires express higher levels of pRUNX2 relative to those cultured on the indicated flat surface (graphs show mean±SD, d=2, r=3, t=17,23, stats by T-test where *=p<0.05). All analysis were performed 24 hours after bacterial inoculation of a seeded MSC culture. Data shows that in co-culture, the pPEA-coated nanowires support MSC growth while preventing biofilm formation.

### Nanowires rescue MSCs from the effects of quorum sensing molecules

Our results show that pPEA-coated nanowires can reduce the adhesion of bacteria in the presence of MSCs. We therefore hypothesised that this surface topography might also protect MSCs from the cytotoxic effects of quorum sensing signalling molecules (QSSMs). To investigate this idea, we seeded MSCs onto pPEA-coated and uncoated flat and nanowire surfaces in the presence of N-(3-oxododecanoyl)homoserine lactone (C12-HSL, Figure 6a), which is a QSSM believed to effect cell viability by disrupting nuclear factor-κB (NFkB) and by inducing apoptosis.^58, 59^ First, we identified a concentration of C12-HSL that could robustly effect MSC viability; 200 μM C12-HSL was selected as this concentration produced a consistently detrimental effect on MSCs. As Supplementary Figures 5 and 6 show, most C12-HSL concentrations tested were highly cytotoxic.

**Figure 6.**
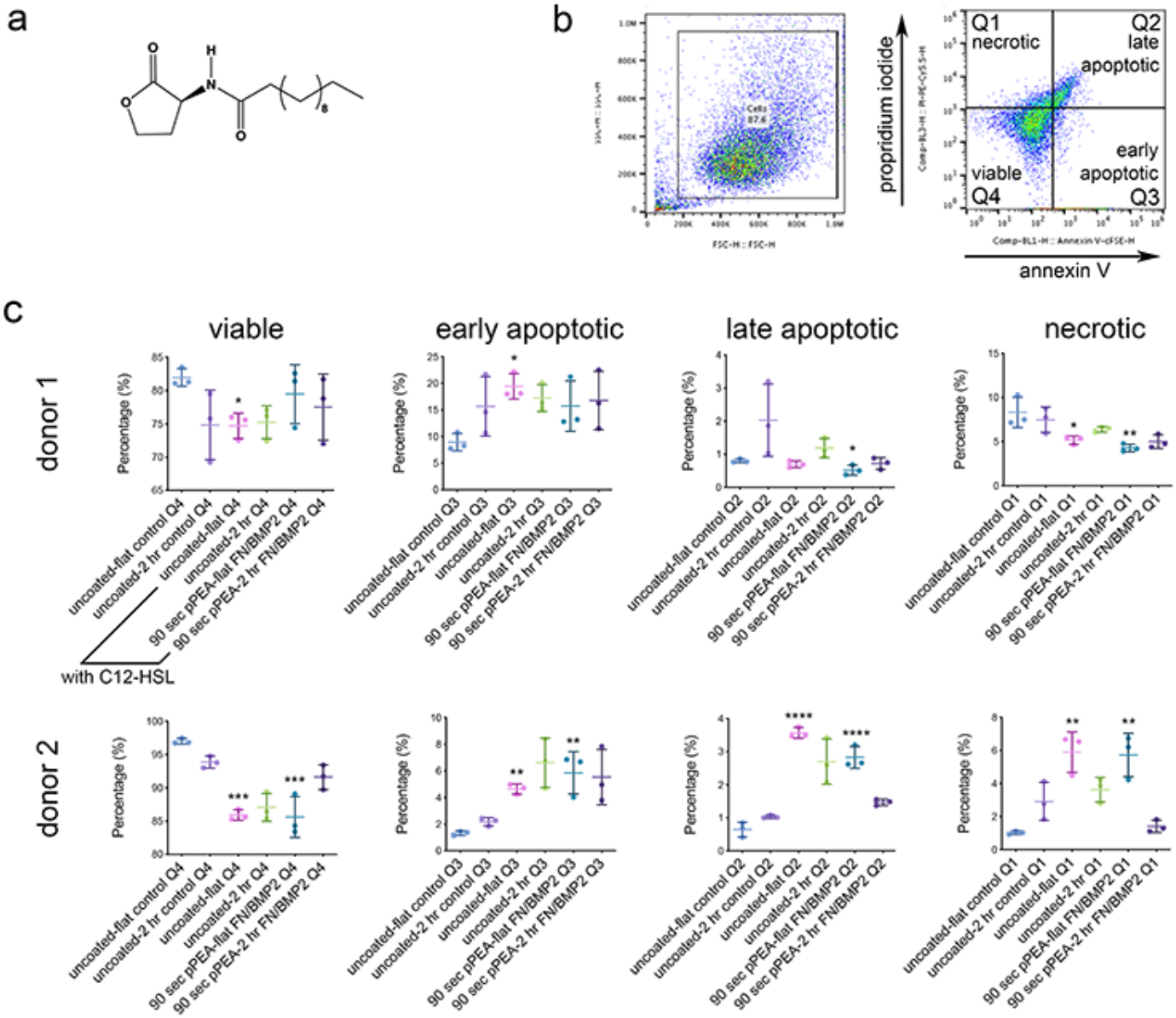
Effects of the quorum-sensing signalling molecule, C12-HSL, on MSCs cultured on uncoated and pPEA-coated nanowires. (a) The structure of C12-HSL. (b) Gating strategy used for flow cytometry analysis, which allowed MSC cells to be separated on the basis of size and viability. (c) Quantification of viable, early apoptotic, late apoptotic and necrotic MSCs after 24 hours of culture on control and test surfaces, with and without C12-HSL (graphs show mean± SD, d=2, r=3, t=1, stats by ANOVA and Tukey test where *=p<0.05, **=p<0.01, ***=p<0.001 and ****=p<0.0001). The results show that nanowire surfaces protect MSCs from the cytotoxic effects of C12-HSL.

To test MSC viability on the different topographies and coatings in the presence of 200 μM C12-HSL, we used annexin V / propidium iodide staining and flow cytometry to analyse MSC apoptotic and necrotic death. The gating strategy we used allowed viable, early apoptotic, late apoptotic, and necrotic phenotypes to be assessed (Figure 6b). No differences in MSC viability were noted in the absence of C12-HSL when MSCs were cultured on uncoated, control flat surfaces and on uncoated nanowire samples; all cells remained highly viable (Figure 6c). However, once 200 μM C12-HSL was added to MSC cultures, apoptosis and necrosis increased in cells cultured on both uncoated and pPEA-coated flat samples (Figure 6c). However, MSC viability was unaffected by C12-HSL when the cells were cultured on uncoated or pPEA-coated 2 hr nanowire surfaces; this was particularly notable with MSCs from donor 2 (Figure 6c). These data indicate that the nanowire topographies help to protect MSCs from cytotoxic QSSMs.

### 3D proof of concept

Trabecular metal and 3D-printed orthopaedic implants are growing in popularity as they can be matched to the modulus of bone and allow bone ingrowth.^60^ Metals, such as titanium and its alloys, are widely used in orthopaedics due to their biocompatibility, resistance to corrosion, and ability to carry load. However, while cortical bone has a Young’s modulus of around ~18 GPa (~10 GPa for trabecular bone), bulk Ti has a modulus of >100 GPa.^61, 62^ This modulus mismatch results in stress shielding because the implant can’t bend with the bone, leading to bone loss through loss of cellular mechanotransduction.^63^ The primary fixation of an implant occurs through bone-implant interdigitation, and is a vitally important process for preventing implant micromotion and aseptic loosening. Porous implants are thus ideal as they can maximise primary fixation.^63, 64^ Secondary fixation occurs at a cellular level and is also vitally important^64^. We therefore hypothesised that our coated nanotopographies can reduce bacterial adhesion and enhance secondary fixation of 3D implant structures.

To investigate this possibility, we used selective laser melting (3D printing) of Ti-6Al-4V ELI (a commonly used orthopaedic Ti alloy) to create porous implants of different strut sizes (300, 600 and 900 μm diameter struts; labelled as D300, D600 and D900, respectively). This 3D printing process uses a laser source to melt Ti-6Al-4V powders (20-63 μm diameter) into 3D shapes; here, the unit cell is a body centred cubic design (Figure 7a). Instron uniaxial compression was then used to calculate Young’s modulus, showing that D900 has a Young’s modulus of ~2 GPa, approaching the natural stiffness of bone (Figure 7a). Further processing of the D900 samples to fabricate the 2 hr nanowires had negligible effects on the Young’s modulus (Figure 7a).

**Figure 7.**
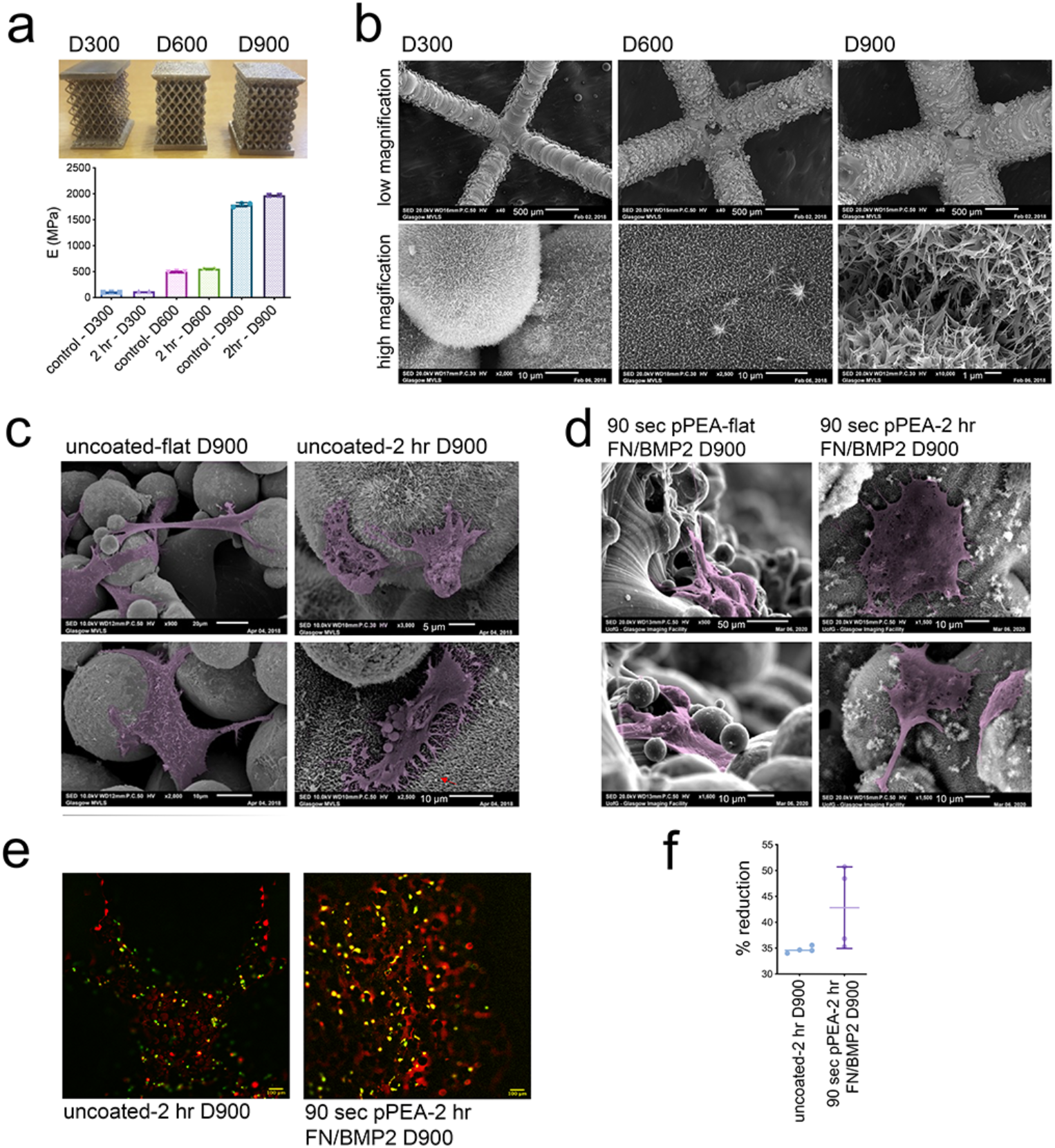
Fabrication of 3D pPEA-coated nanowires and MSC responses to 3D surface. (a) The 3D printed implants, showing their different strut diameters (D300, D600 and D900), and (lower panel) the measured Young’s modulus for each 3D shape (graphs show mean±SD, n=2,3). (b) SEM of 2 hr nanowires on the 3D struts of each implant. (c) SEM of MSCs cultured for 3 days on uncoated D900 samples, showing reduced spreading on 2 hr nanowires compared to flat control samples. (d) SEM of MSCs cultured for 3 days on a 90 sec pPEA-coated D900 sample, showing enhanced cell spreading on the 2 hr nanowires compared to the flat control sample. (e) Fluorescent staining of actin (red) and osteopontin (green) in MSCs cultured on an uncoated 2-hr D900 sample and a 90 sec pPEA-coated FN/BMP2, 2-hr D900 sample for 21 days. The pPEA FN/BMP2 coating increased osteopontin expression, relative to the control. (f) Quantification of Alamar blue staining in MSCS after 14 days of culture, indicating that the pPEA coating has no detrimental effect on MSC viability cultured in 3D (graph shows mean±SD, d=1, r=4, t=3, stats by T-test >0.05).

When the D900 samples were analysed by SEM following the addition of the nanowires, it revealed the fused-particle nature of the implants and that the struts have a good topographical coating of nanowires (Figure 7b). SEM also showed that adding nanowires to D900 reduced MSC spreading on this surface, relative to uncoated, flat D900 (Figure 7c). However, MSC spreading was rescued by the use of the pPEA FN/BMP2 coating (Figure 7d). Immunofluorescent staining for the bone marker osteopontin also confirmed our earlier 2D observations (Figure 2) that the pPEA FN/BMP2 coating enhances osteogenesis (Figure 7e) when MSCs are cultured on nanowire surfaces. The pPEA FN/BMP2 coating also appeared to have no cytotoxic effect when applied to the 3D implants, as assessed by Alamar blue staining (Figure 7f). Finally, SEM also confirmed the antibacterial effect of the pPEA-coated nanowires in 3D, as observed in 2D (Figure 4), and demonstrated reduced biofilm formation on pPEA-coated nanowires compared to that seen in uncoated flat controls (Figure 8a). Again, it was noted that while *P. aeruginosa* had numerous flagella on the uncoated flat D900 controls, no flagella were observed on the pPEA-coated 2 hr nanowire D900 samples and there was evidence of bacterial rupture (Figure 8b).

**Figure 8.**
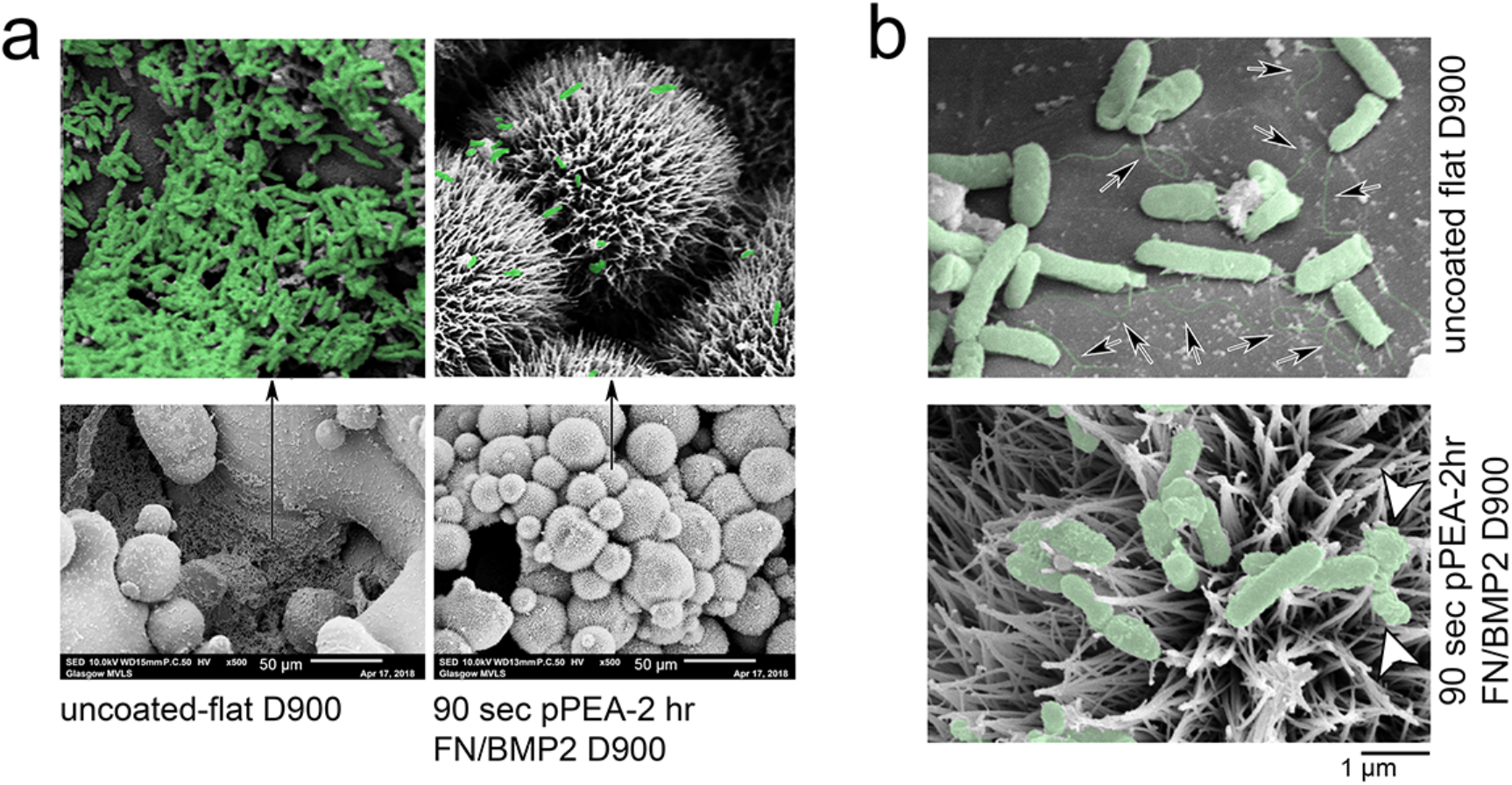
*P. aeruginosa* response to 3D pPEA-coated nanowires. (a) SEM of *P. aeruginosa* cultured on the D900 uncoated flat control and on the 90 sec pPEA-2 hr FN/BMP2 coated D900 surface. In the uncoated flat samples, large areas of confluent biofilms were observed. By contrast, only small areas of bacterial accumulation were seen on the pPEA-coated nanowires and much of the implant was clear of bacteria. (b) Flagella were observed in all bacterial cells on D900 uncoated flat control samples (arrows) but were absent in bacterial cells observed on 90 sec pPEA-2 hr FN/BMP2 coated D900. Ruptured bacterial cells were also observed on the nanowire surfaces (arrowheads).

## Summary

In this study, we set out to show that by exploiting both the cell adhesion and growth factor binding properties of the ECM using PEA coatings that we could induce osteogenesis of MSCs on antibacterial topographies. Our findings confirm earlier reports that high aspect ratio nanowires have bactericidal activity^45, 46, 48^ (Figure 4b) and can be fabricated into relevant biomaterials, such as titanium.^47^ These viability effects occur together with reduced adhesion, producing higher levels of planktonic culture (Figure 5 b,c) with sessile bacterium either ruptured (Figures 5c and 8b) or trapped onto the nanotopography’s surface features (Figures 4a and 8b). The topographies also resulted in the loss of flagella in the *P. aeruginosa* cultures (Figures 4a and 8b), indicating cell stress.^55^

We also investigated here whether we could use nanoscale pPEA coatings to present growth factors, such as BMP2, in synergy with integrins in solid phase through open-conformation FN. PEA provides a means of delivering growth factors topically and at low concentration to reduce their systemic effects^19^. It has also been used in veterinary trials to treat critical sized, non-healing, fractures, where allograft was coated in PEA with FN/BMP2.^53^ Here, to preserve the sharpness of the nanowires, we used a very thin PEA coating of ~40 nm for our optimal, 90-sec coating, rather than the ~300 nm coating used in the veterinary trial graft. This very thin film led to a heterogeneous surface covering of PEA, with Ti detectable by XPS, as well as PEA (Figure 1f); it also led to the formation of both globular and fibrillar FN (Figure 3d). It is thus interesting that MSCs responded better, in terms of bone marker gene expression and FN nanonetwork formation, to the more homogeneous PEA coating produced by the 3min-coating (of 70 nm thickness). It is possible that cells are more used to heterogeneous ECM environments rather than to highly conformal, optimised, experimental conditions. Our findings also show that PEA-coated nanowires enhanced MSC osteogenic responses, relative to those produced by MSCs cultured on uncoated nanowires^47^ or on integrin mimetic coated nanowires,^49^ likely because of the synergy between RGD and BMP2.^19, 22^ We note that the PEA/FN/BMP2 interface is very stable and that over 2 weeks, <8% of absorbed BMP2 is released.^19, 53^ Our nanowire data indicate that these structures absorb more FN than does a planar surfaces, perhaps as a function of surface area, and that more of the RGD-containing FN region is available to cells when a surface’s features are pPEA coated (Figure 3g). By contrast, the availability of the heparin binding site and BMP2 absorption levels remained unchanged by pPEA coating. However, between 70-80% of the BMP2 was absorbed onto the FN, and we propose that this solid-state BMP2 coupled with increased RGD availability can drive synergistic signalling between RGD and BMP2, as previously reported.^19, 53, 54^

A key finding of this study is the reduction in bacterial adhesion seen when *P. aeruginosa* is cocultured with MSCs on pPEA-coated nanowires. While biofilms formed swiftly across the surface of flat Ti, very little bacterial adhesion was observed on the nanowires, in both the presence or absence of the pPEA FN/BMP2 coating (Figure 5a, b). However, the pPEA FN/BMP2 coating did increase MSC adhesion and pRUNX2 expression (Figure 5d,f) highlighting its osteoinductive potential, even in MSCs co-cultured with pathogenic bacteria. We recognise that our use of antibiotics to reduce bacterial growth is a limitation of this study, as is the short term nature of the co-culture due to the bacterial killing of MSCs. However, in the clinic, antibiotic use is common, especially in revision surgery.^65^ Furthermore, low bacterial adhesion and good MSC adhesion both help regenerative cells win the ‘race for the surface’ that is considered to be very important for preventing implant infection.^66^

We also demonstrate that the well-known QSSL virulence molecule, C12-HSL, has reduced cytotoxic effects on MSCs cultured on the nanowire topography, particularly when coated with pPEA FN/BMP2 (Figure 6c). This allows us to hypothesise that enhanced MSC-FN attachment and synergistic BMP2 signalling protects MSC viability. It is also noteworthy that while the solid-phase delivery of BMP2 reduced *P. aeruginosa* amino acid metabolism (Figure 4d), its soluble delivery caused increased pathogen metabolism (Supplementary Figure 3).

Finally, as proof of concept, we generated 3D-printed Ti alloy implants coated with nanowires and with pPEA FN/BMP2, thereby generating porous implants with a stiffness that more closely resembles that of bone than Ti implants. When the pPEA FN/BMP2 coating was applied to this 3D structure, it again enhanced MSC adhesion and increased the expression of the bone marker osteopontin (Figure 7c-e), while the inclusion of nanowires on this 3D structure reduced biofilm formation (Figure 8a, b).

## Conclusions

The use of high aspect ratio nanotopographies to reduce infection is an area of growing interest. In this study, we show that to realise the potential of such coatings on orthopaedic implants, when infection control is required, consideration must also be given to supporting the bone-forming MSCs, as these cells are the primary regenerative target of the implant. Our findings show that by using a very thin pPEA coatings on an implant, it is possible to organise FN and BMP2 in a biomimetic manner to improve MSC attachment to the surface of an implant and their osteospecific differentiation. This pPEA coating when used on a nanowire topography can also protect MSCs from QSSL virulence factors and reduce bacterial adhesion. These protective qualities and enhanced MSC spreading both help MSCs to win the race to colonise the implant’s surface. The nanotechnologies presented here are facile and can be fabricated in 3D using emerging metal implant additive manufacturing approaches.

## Materials and methods

### Nanowire fabrication and PEA coating

Ti samples were prepared as 0.9 mm thick, 10 mm dimeter disks in ASTM grade one Ti (Baoji HeQiang Titanium Industry Co.). The method of nanowires production has been described in detail in Tsimbouri et al. (2016).

Ethyl acrylate (EA) was polymerized using plasma polymerisation to produce PEA coatings as described in Alba-Perez et al. (2020). PEA was applied on the Ti surfaces for 90 seconds and 3 minutes at 100 Watts. Human FN (R&D Systems) was adsorbed on the top of the surfaces by immersing them in 200 μl of a 20 μg/ml human plasma FN/PBS solution for 1 hr. Samples were washed once with PBS, blocked in 1 % BSA for 30 mins and then washed once with 1x PBS. Growth factor (BMP2) (R&D Systems) was then absorbed in the same way at a concentration of 100 ng/ml.

### Characterization

The surface macroscopic topography was analyzed using atomic force microscopy (AFM). Images were obtained on flat and nanowire samples pre- and post-coating with PEA. AFM experiments were performed using a Multimode AFM equipped with NanoScope IIIa controller from Veeco (Manchester, UK) operating in tapping mode in air; NanoSscope 5.30r2 software was used. Average roughness (*R_a_*) values and standard deviation are reported.

The sessile drop contact angles were measured on the surfaces as an indication of the wettability of the substrates. Droplets of deionized water (volume ≈ 3 μL) were deposited at different positions on the surfaces (Optical Tensiometer Theta, Biolin Scientific). The results are presented as the mean contact angle of at least eight droplet measurements per condition.

X-ray photoelectron spectroscopy (XPS) was used to analyse the chemical components of the metal alloys pre- and post-plasma PEA coating. The spectral measurements were obtained at the National EPSRC XPS Users’ Service (NEXUS) at Newcastle University, an EPSRC Mid-Range Facility. XPS was performed using a K-Alpha apparatus (Thermo Fisher Scientific), with a micro-focused monochromatic Al Kα source (X-ray energy = 1486.6 eV) at a voltage of 12 kV, current of 3 mA, power of 36 W, and spot size of 400 μm × 800 μm. Spectral analysis and curve fitting were performed using CasaXPS software version 2.3.16.

### Protein coating quantification

**A.** FN adsorption: The protein adsorption properties of the FN coating were analysed by performing a bicinchoninic acid (BCA) protein assay by following the manufacturers protocol (Thermo Fisher Scientific). **B.** BMP2 adsorption: The amount of BMP2 bound to the surfaces was quantified by indirect enzyme-linked immunosorbent assay (ELISA) using DuoSet human BMP2 kits’ (R&D Systems). After the samples were coated with BMP2, the coating solutions and the first wash solutions were collected for analysis. The amount of BMP2 bound to the samples was calculated as the difference between the total BMP2 used for coating and the amount of BMP2 in the collected solution. **C.** Integrin- and heparin-domain viability: P5F3 and HFN7.1 antibodies were used to assess the availability of FN’s heparin-binding domain and cell-binding domain, respectively. Primary mouse anti-human antibodies P5F3 (Santa Cruz) and HFN7.1 (AB_528244, DSHB Hybridoma Product HFN7.1) were added onto the coated surfaces and were incubated for 1 hr at RT. The samples were then washed three times with PBS/0.5% Tween 20 and subsequently incubated with an anti-mouse IgG-HRP-conjugated 2ry antibody (Thermo Fisher Scientific) for 1 hr. Following two washes, the samples were incubated in 200 μl of substrate solution (tetramethylbenzidine, TMB solution) (R&D systems) per sample for 20 min. 100 μl of stop solution (Sulfuric acid) (R&D systems) was added. A microplate reader was used to determine optical density at 450 and 570 nm.

### Cell culture

Stro-1^+^ selected primary human MSCs from adult human bone marrow, obtained with informed consent, were provided by Southampton General Hospital. MSCs were cultured in basal media, (Dulbecco’s modified essential medium (DMEM; Sigma) supplemented with 10% fetal bovine serum (FBS; Sigma), 1% (v/v) L glutamine (200 mM, Gibco), 1% sodium pyruvate (11 mg/mL, Sigma), 1% MEM NEAA (amino acids, Gibco), and 2% antibiotics (6.74 U/mL penicillin-streptomycin, 0.2 μg/mL fungizone; Sigma). MSCs were cultured in an incubator set at 37 °C with 5% CO_2_ environment and sub-cultured to passage 2-3 before use. Culture media was changed every 3 days. Seeding on coated Ti was done in serum-free medium in 24-well plates at 10^4^ cells/disc. After 2 hr, the media were changed to low serum growth medium (1% FBS).

### Quantitative polymerase chain reaction with reverse transcription

Total RNA was extracted at relevant time points using a Qiagen RNeasy Micro kit and protocol. An RNA extraction kit (RNeasy extraction microKit, Qiagen) was used to purify RNA. The concentration of purified RNA was measured by spectrophotometer (Nanodrop 2000c, Thermo Fisher Scientific). cDNA was then synthesized using the QuantiTect Reverse Transcription Kit (Qiagen). Forward and reverse primers for qRT-PCR are shown in Table 1. GAPDH, a house-keeping gene, was used as internal control of the analysis. SYBR Green dye was used to target synthesized cDNA (Quantifast SYBR Green I, Qiagen). Real-time PCR was then performed (7500 Real Time PCR system, Applied Biosystem, USA). The 2^−ΔΔCT^ method was used for interpretation.^67^

**Table 1:**
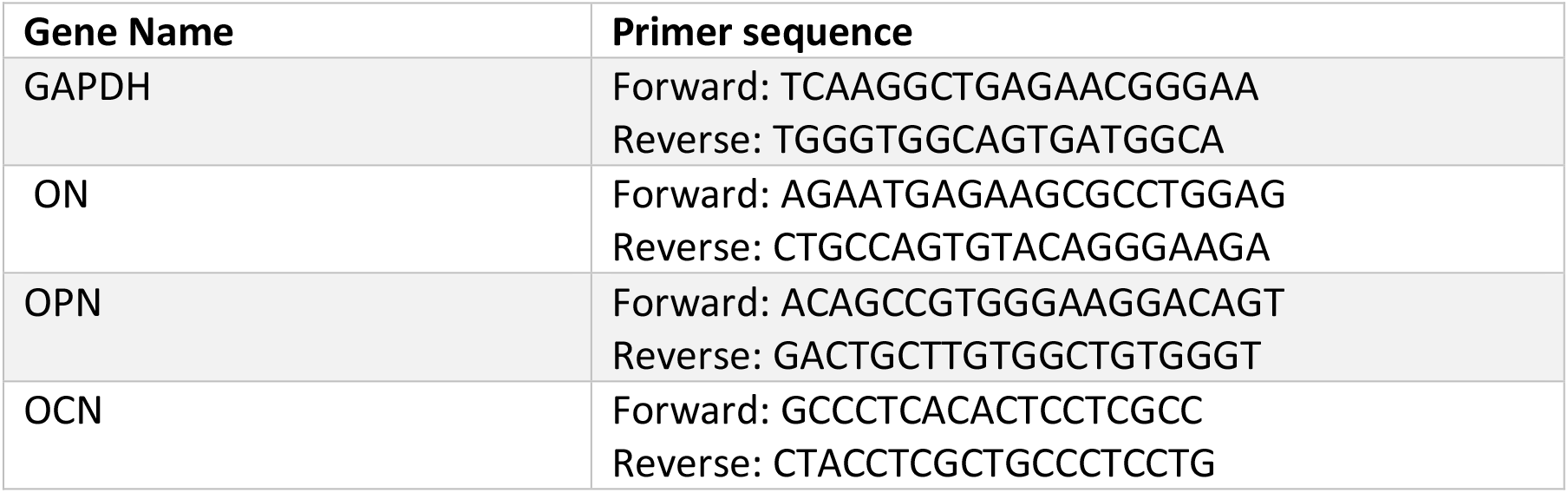
Primers used for qRT-PCR.

### Alizarin red S and calcein blue

Mineral production was assayed using Alizarin red and calcein blue staining. For Alizarin red staining, after 35 days in culture, cells cultured on flat and nanowires surfaces were washed with 1x PBS and fixed with 4% v/v formaldehyde 1x PBS for 15 min at 37 °C. 1 ml/well Alizarin red stain solution (40 mM) was added and incubated at RT for 30 minutes. Excess dye was removed, and cells were carefully washed 5x with deionized water. Calcium deposits were visualised using a light microscope (Leica, Germany).

For calcein blue staining, 10 mg of calcein blue (Sigma) were dissolved in 0.25 ml of KOH (1 M) and 9.75 ml of distilled water were added to make up a 3.1 × 10^−3^ M calcein blue solution. 5 μl of calcein blue solution were added to the culture medium (final concentration of calcein blue: 3.1 X10^−5^ M) (1:100) was added to the culture medium 1 h before cell fixation. The cells were then washed with PBS 3x and fixed with 4% formaldehyde in PBS for 15 min at 37 °C. After fixation, the specimens were washed with PBS three times and dried. Stained cells were imaged using a Zeiss immunofluorescence microscope. Images were processed using ImageJ imaging software, and images that were to be compared against one another were taken with the same exposure parameters.

### Bacteria culture

*P. aeruginosa* ATCC 27853 was grown aerobically overnight in 10 mL DMEM in a 37 °C shaker incubator set at 220 rpm. The bacterial suspension was then diluted in DMEM to OD_600_= 0.1 and further incubated until mid-exponential phase was reached. At this time, bacterial cells were harvested by centrifugation (7 min, 5000 xg), washed twice in PBS buffer, and suspended in DMEM OD_600_= 0.3 (approx. 10^8^ CFU/mL). 10^4^ CFU/ml were used for culture.

### Bacteria imaging and quantification

To evaluate viable bacterial cell numbers based on the ATP present, the nanowire surfaces and flat controls were placed into a 24 well plate and submerged in the bacterial suspension. Plates were incubated for 18 hr at 37 °C under static conditions. After incubation, 100 μl of each supernatant were transferred into new 96 well plates and equal volume of BacTiter-Glo™ Reagent (Promega) was added per well. The plates were read within 5 minutes of adding the reagent using a luminescence plate reader (CLARIOstar, BMG LABTECH, Germany).

Adhesion of bacteria on different types of material was observed using an inverted Zeiss Axiovert fluorescence microscope using live/dead staining (LIVE/DEAD^®^ BacLight™ bacterial viability kit, Molecular Probes); the green and red intensities were used to determine the threshold between live and dead bacteria, respectively.

### Metabolomics

*P. aeruginosa* was cultured on different Ti surfaces for 24 hr. After incubation, each Ti disc was transferred into a bijou tube and kept on ice. A 1 ml: 1g mixture of a chloroform, ethanol and ddH_2_0 (CEW, ratio 1:3:1) buffer and acid washed glass beads 0.1 mm was produced, and 1 ml was added to each bijou. Samples were placed on a cell disrupter (Disrupter Genie bead beater, Scientific industries Inc., New York, USA) operating at a speed of 3000 rpm, 3x for 30 sec. The entire liquid/bead mixture from each bijou was transferred to microcentrifuge tubes and centrifuged at 10,000 rpm for 3 minutes to remove the beads. The supernatant was subsequently transferred to a fresh microcentrifuge tube. A pool sample was produced by adding 20 μL of each sample to a separate tube. Each metabolomic sample contained a minimum of 200 μL of cell extract. Samples were then stored at −80°C until metabolomic analysis could be performed. Hydrophilic interaction liquid chromatography-mass spectrometry was performed (Dionex, UltiMate 3000 RSLC system, Thermo Fisher Scientific, Hemel Hempstead, UK) using a ZIC-pHILIC column (150 mm × 4.6 mm, 5 μm particle size, Merck Sequant). The data sets were processed using XCMS (peak picking), MzMatch (filter and grouping), and IDEOM (post processing filtering and identification). Metaboanalyst was used to generate heatmaps and principle component analysis (PCA).

### MSCs and *P. aeruginosa* co-culture

MSCs were cultured on the different Ti substrates and incubated for 24 hr before adding the bacterial culture at the concertation 10^3^ CFU/ml with 0.3% of Ab (Supplementary Figure 5). After 24 hr incubation the samples were rinsed with PBS for SEM and IF images.

### Scanning electron microscopy (SEM)

After 24 hr of co-culture, the samples were fixed in 1.5% glutaraldehyde for MSCs for 1 hr at RT or 2.5 % glutaraldehyde for bacteria buffer overnight at 4°C and then rinsed in 0.1 M sodium cacodylate. Samples were post fixed in 1% osmium tetroxide for 1 hr at RT then washed 3x with distilled H_2_O for 10 min. The samples were then stained by 0.5% uranyl acetate for 1 hr in the dark. For dehydration, a 30-100% ethanol series was applied. A hexamethyldisiloxane step was conducted prior to sputter coating (20 nm gold/palladium). Samples were viewed on a JEOL IT100 SEM, running at 10-20 Kv, on Secondary Electron Detector (SED) mode. Tiff images were captured using JEOL Intouch Scope software version 1.03.

### Immunofluorescence (IF)

For protein detection by immunofluorescence, two independent experiments were set up with 3 technical replicates per condition. Briefly, cells were fixed in 4% formaldehyde fixative at 37 °C for 15 min and permeabilized at 4 °C for 5 min. The samples were blocked with 1% BSA/PBS at 37 °C for 5 min and stained with the appropriate primary antibodies for pRUNX-2 or OPN (1:50 in 1% BSA/PBS, Autogen Bioclear). Actin filaments were stained with phalloidin (1:100, Invitrogen). The samples were washed in 1 × PBS/0.5% Tween-20 (3 × 5 min at RT) and a secondary, biotin-conjugated antibody (1:50, horse monoclonal anti-mouse (IgG), Vector Laboratories) was added for 1 hr at 37 °C. The samples were washed, and fluorescently conjugated streptavidin was added (1:50, Vector Laboratories) at 4 °C for 30 min, before the samples were given a final wash and mounted in Vectashield mounting medium (Vector Laboratories) containing DAPI to stain the nucleus. Visualisation was via a fluorescence microscope (Zeiss Axiovert 200 M, 10x magnification, NA 0.5). Comparisons of staining intensity between substrates were analysed by Image J software version 1.42q.

### MSCs culture with C12-HSL

MTT (3-(4,5-dimethylthiazol-2-yl)-2,5-diphenyltetrazolium bromide, Sigma) assay was used to evaluate the metabolic activity of cells in the presence of quorum sensing molecule C12-HSL. After 24 hr incubation with C12-HSL, 10 μl of MTT dye solution (5 mg/ml MTT in PBS, pH 7.4) was added to each well of a 96-well plate and incubated for 2 hr. After the incubation, formazan crystals were solubilised with 200 μl DMSO. The absorbance of each well at 550 nm was read on a Dynatech MR7000 microplate reader. A parallel experiment with live/dead staining (LIVE/DEAD^®^ Viability/Cytotoxicity Kit for mammalian cells, Molecular Probes) was performed.

### Flow cytometry

MSCs were incubated with 200 μM C12-HSL for 24 hr and then stained with an antibody against Annexin V to detect apoptosis. After incubation, the cells were harvested by adding 200 μl of trypsin and washed with PBS and another wash was applied using 1x Annexin V binding buffer (10x concentrate composed of 0.1 M Hepes (pH 7.4), 1.4 M NaCl, and 25 mM CaCl2 in PBS). The cells were incubated with 5 μl of Annexin V staining for 15 mins/ dark/ RT. 500 μl of binding buffer were added to wash the cells and samples were centrifuged at 419 xg for 4 mins. Fresh 200 μl of binding buffer were added and 1 μl of PI before flow cytometry was performed (Thermo Fisher Scientific). The excitation/emission for Annexin V is 494/518, and 535/617 for PI. The cells were gated, and standard compensation was applied (Figure 6b and Supplementary figure 5a) (Attune NxT, life technologies).

JC-1 Dye was used for mitochondrial membrane potential. The MSCs were incubated with 200 μM C12-HSL. After the 24h incubation the supernatant was removed and fresh media containing 2 μM JC-1 were added to the cells; for the control tube, 50 μM CCCP was added and incubated at 37°C, 5% CO_2_, for 30 mins. The cells were washed once with PBS, trypsinized and pelleted by centrifugation at 419 xg for 4 mins. The cells were resuspended in fresh PBS and analysed using a flow cytometer with 488 nm excitation using emission filters appropriate for Alexa Fluor 488 dye and R-phycoerythrin. The cells were gated, excluding debris using the CCCP-treated sample, and standard compensation was applied (Supplementary figure 4d).

### 3D-Ti lattice production

A body centred cubic (BCC) design of Ti-6AI-4V was manufactured using the SLM system SLM-250HL. The argon atomised Ti-6Al-4V powder that was used in the SLM process features extra low interstitials (ELI). The samples were produced at the Singapore Centre for 3D Printing, School of Mechanical and Aerospace Engineering, Nanyang Technological University, Singapore. Three different Ti lattices with the following strut diameters D300- D600- D900 μm were produced. The TiO_2_ nanowires were formed on the 3D Ti lattices using the alkaline hydrothermal method, as mentioned before.

### Compression test

The compression on the micro lattices was performed at the quasistatic rate 0.001 s^−1^ in an INSTRON machine, with the deformation history recorded by a JAI camera.

### AlamarBlue assay

Stro^+1^ MSCs were seeded in 3D Ti lattices at a density of 5 ×10^4^ cells/mL for 3 weeks. Samples were washed with warm 1× PBS. 10% (v/v) of AlamarBlue resazurin (Bio-Rad, Watford, UK) was diluted in DMEM. After incubation at 37 °C and 5% CO2 for 6 h, the supernatant containing the taken up AlamarBlue was pipetted and transferred into 96-well plates. A microplate reader (Clariostar, BMG Labtech, Germany) was used to detect light absorbance at wavelengths of 570 and 600 nm.

### Statistical Analysis

All experimental results were interpolated and analysed using GraphPad Prism (GraphPad Software Inc.). Means and standard deviations were calculated, and data were analysed by t test, one- or two-way analysis of variance (ANOVA) test with Tukey’s or Dunnett’s multiple comparison post-test. All results are shown in mean ± standard deviation with 95%, 99%, and 99.9% of accuracy (* *P* ≤ 0.5, ** *P* ≤ 0.01, *** *P* ≤ 0.001, **** *P* ≤ 0.0001). Replicate details for each experiment are shown in Table 2.

**Table 2:** Details of sample replicates and statistical tests used in figures.

| Figure | Sub figure | Name | Statistical analysis | Donors | Material replicates | Technical replicates |
| --- | --- | --- | --- | --- | --- | --- |
| 1 | c | AFM (Ra) | - | - | 4 | 2 |
| 1 | d | WCA | - | - | 4 | 2 |
| 2 | A, b | qPCR | One-way ANOVA | 1 | 4 | 2 |
| 2 | d | Calcein Blue | T-test | 1 | 4 | 15 |
| 3 | b | ELISA | One-way ANOVA | - | 3 | 1 |
| 3 | f | Coating thickness | One-way ANOVA | - | 4 | 1 |
| 3 | g | ELISA | One-way ANOVA | - | 3 | 2,3 |
| 4 | b | Live/Dead bacteria | T-test | - | 3 | 12 |
| 4 | c | ATP release | T-test | - | 4 | 3 |
| 4 | d,e | Metabolomics | - | - | 4 | 1 |
| 5 | e,f | Co-culture | T-test | 2 | 3 | 17,23 |
| 6 | c | Apoptosis (on Ti) | One-way ANOVA | 2 | 3 | 1 |
| 7 | a | Compression test | - | - | 2,3 | 1 |
| 7 | f | AlamarBlue | T-test | 1 | 4 | 3 |
| S2 | - | Metabolomics | - | - | 4 | 1 |
| S4 | a | MTT | One-way ANOVA | 1 | 4 | 3 |
| S4 | c | Live/dead mammalian cells | One-way ANOVA | 1 | 4 | 12 |
| S4 | d | JC-1 | One-way ANOVA | 1 | 4 | 1 |
| S5 | b | Apoptosis (on glass) | One-way ANOVA | 3 | 4 | 1 |

## Acknowledgements

L.A.D. was supported by a scholarship from Jeddah University and the Saudi Arabian Government. The work was also supported by grants from EPSRC (EP/K034898/1) and MRC (MR/S010343/1). We thank Carol-Anne Smith and Marcus Eales for laboratory support and Margaret Mullin for help with microscopy.

## Author Declaration

The authors declare no competing financial interests.

## Author Contributions

LAD, MPT, PL, GR, AN, BS, MJD, MS-S, ROCO conceived the experiments. LAD, MPT, MG, VLH PC, VJ, YX, PL performed the experiments. MG, VLH, PC, VJ, KB, JW, MRS, RMDM, PL, AN, GR, BS, ROCO, MS-S provided materials and expertise. LAD, PMT, MJD, MRS, VJ, MJD, PL analysed the data. MS-S, RMDM, ROCO provided critique and context for the data. LAD, MPT, MJD wrote the manuscript. LAD, MJD prepared the figures. All authors read and commented on the manuscript.

## Supplementary data

**Supplementary figure 1.**
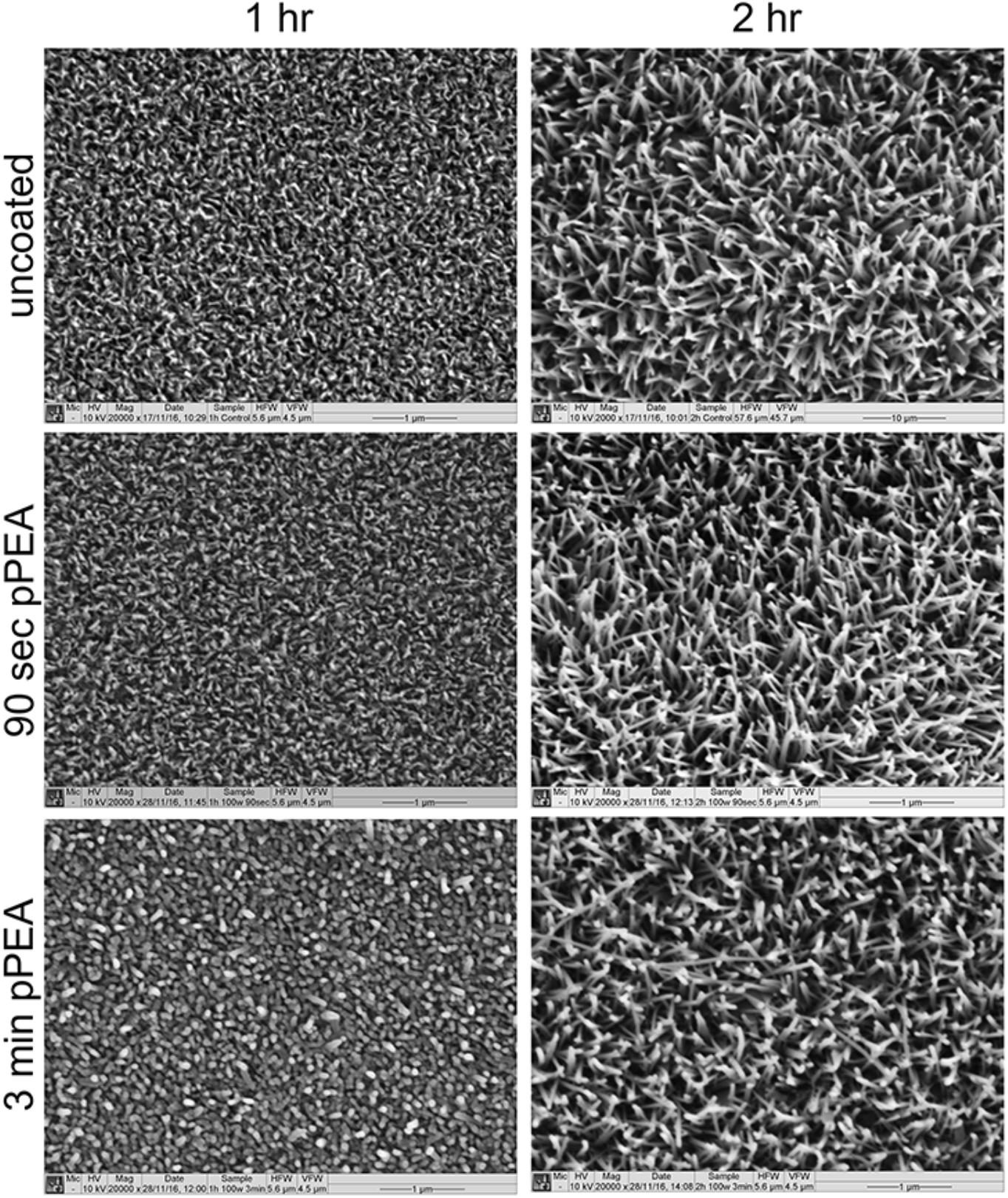
SEM imaging of Ti nanowire surfaces with and without PEA coating. The images show that height of the nanofeatures increase proportionally with anodisation time and that the thin PEA coatings to not effect nanowire morphology.

**Supplementary figure 2.**
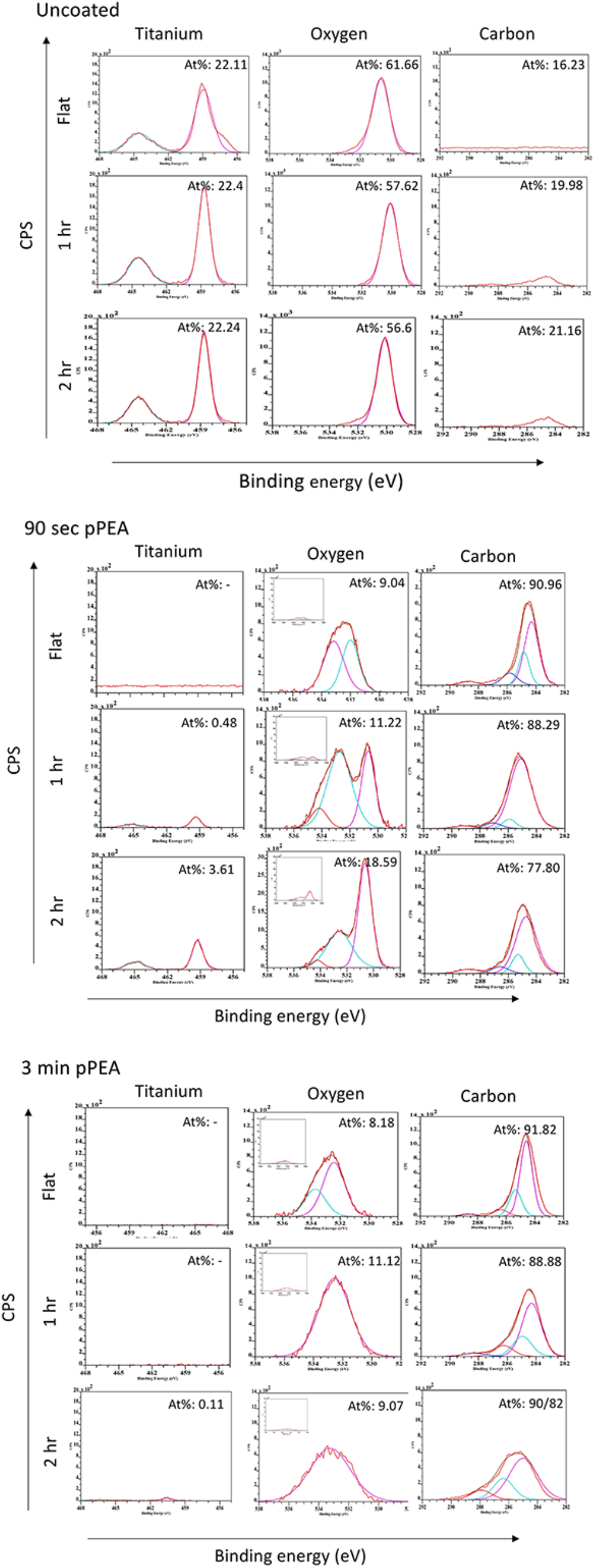
Enlargements of XPS images from Figure 1f.

**Supplementary figure 3.**
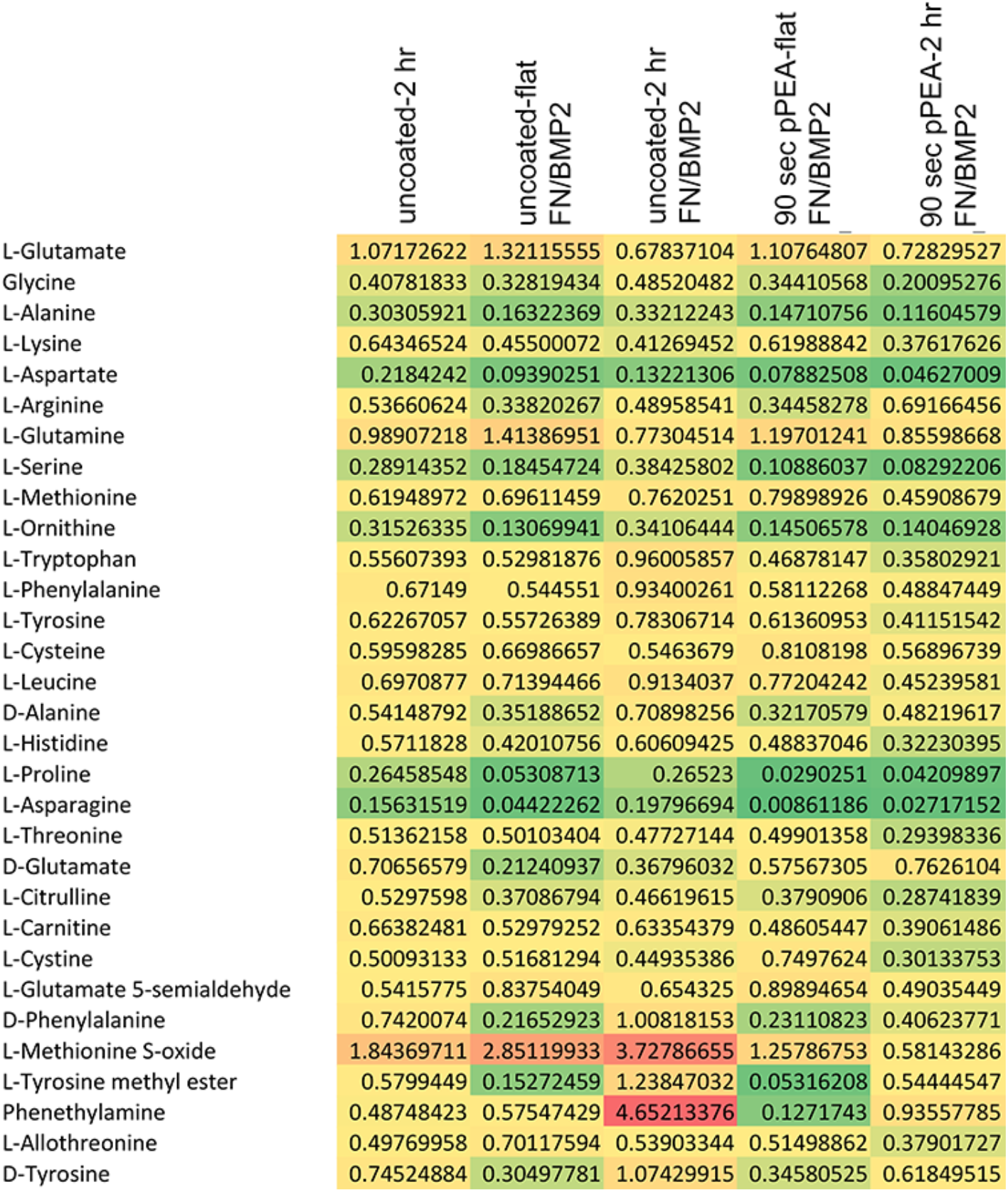
*Untargeted metabolomics data for P. aeruginosa* amino acid metabolism demonstrating that addition of soluble FN and BMP2 can cause increased metabolism while solid state, PEA presentation, reduces amino acid metabolism. Results represent the mean where green = down-regulation, yellow = no change and red=upregulation (r=4, t=1).

**Supplementary figure 4.**
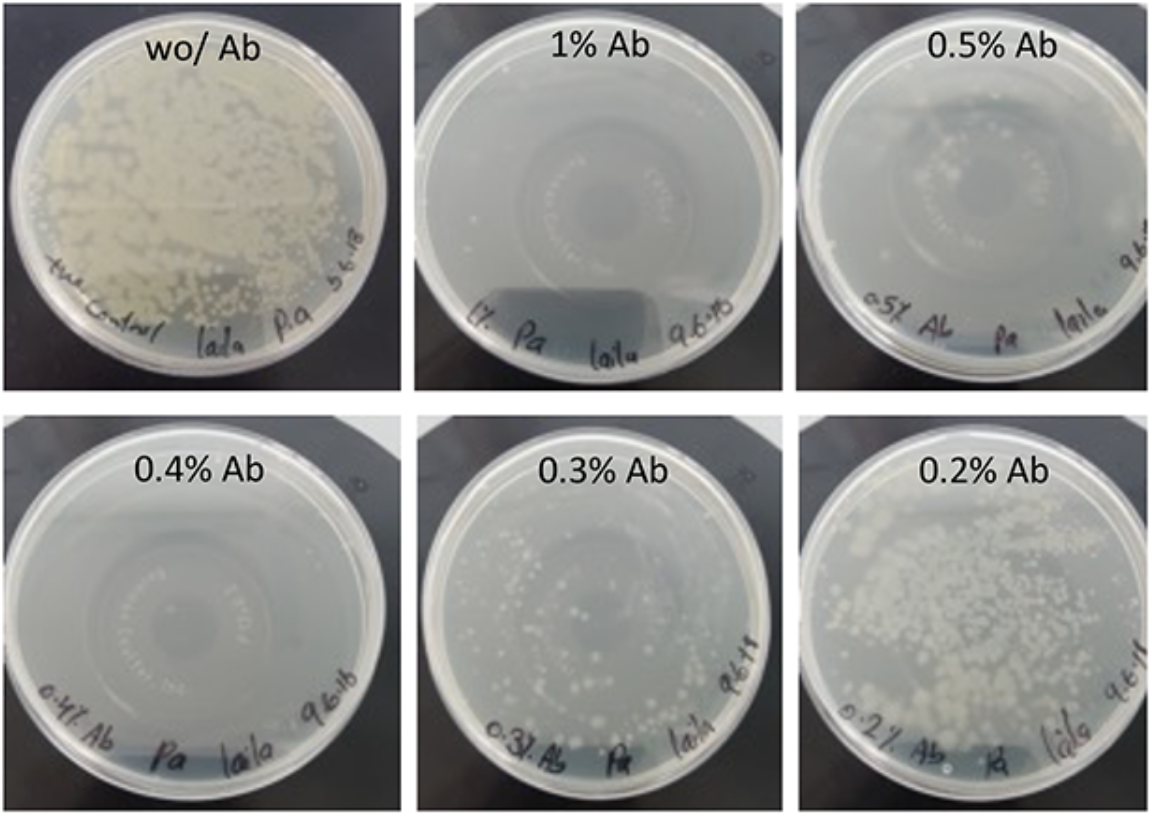
P. aeruginosa growth with different concentrations of antibiotic after 24 hours of plating 50 μl from 10^3^ CFU/ml culture. 0.3% was selected as a concentration that slowed the bacterial growth but that did not kill them.

**Supplementary figure 5.**
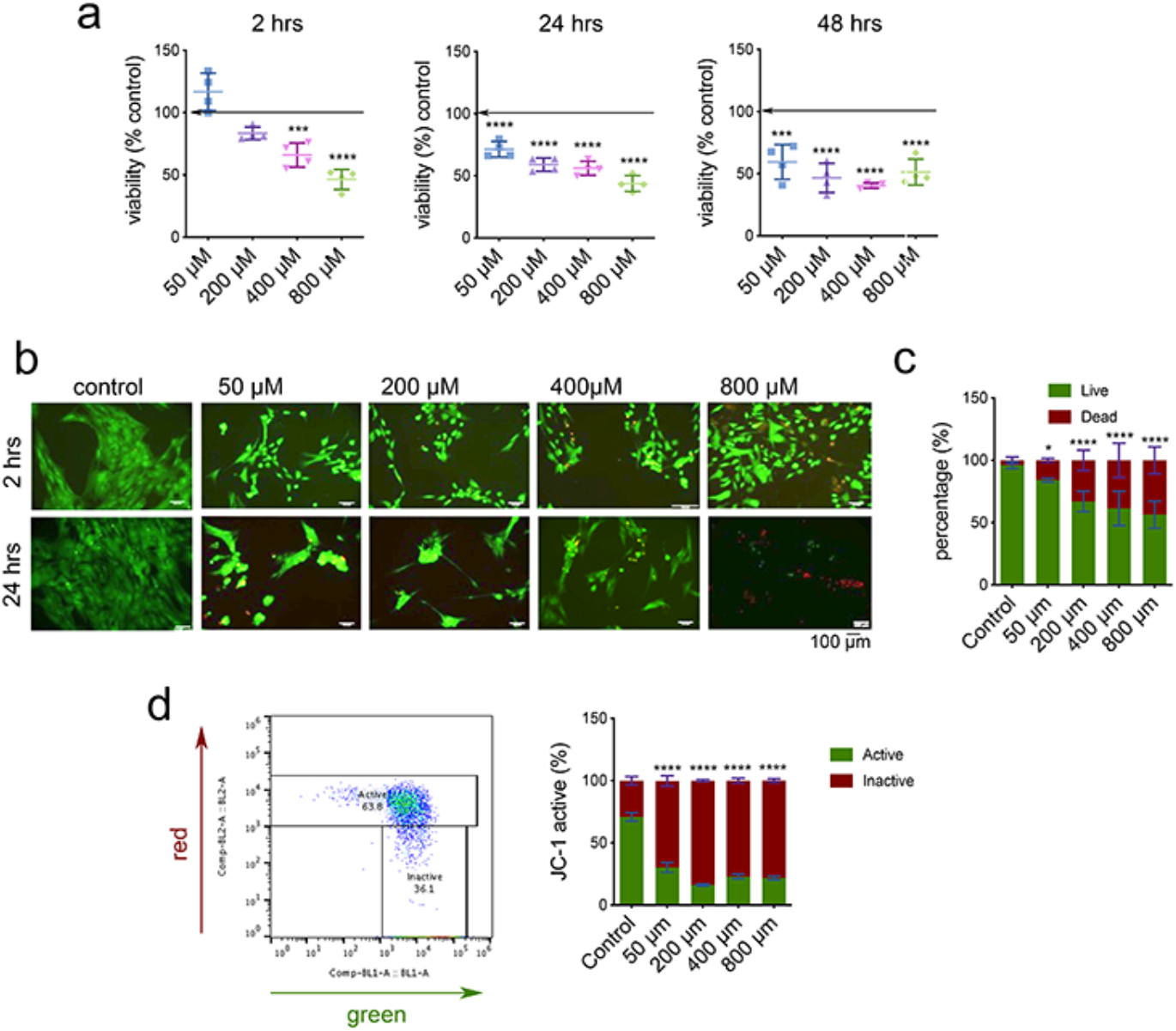
MSC viability assessment with different concentrations of C12-HSL. (a) MTT metabolic assay at 2, 24 and 48 hours of culture with C12-HSL (graphs show mean±SD, d=1, r=4, t=3, stats by ANOVA and Tukey test where ***=p<0.001 and ****=p<0.0001). (b&c) live/dead viability quantification from use of calcein AM and ethidium homodimer fluorescence stain (graphs show mean±SD, d=1, r=4, t=9, stats by ANOVA and Tukey test where *=p<0.05 and ****=p<0.0001). (d) JC-1 staining for mitochondrial activity showing gating strategy and % mitochondrial activity with increasing C12-HSL concentration (graphs show mean±SD, d=1, r=3, t=1, stats by ANOVA and Tukey test where ****=p<0.0001).

**Supplementary figure 6.**
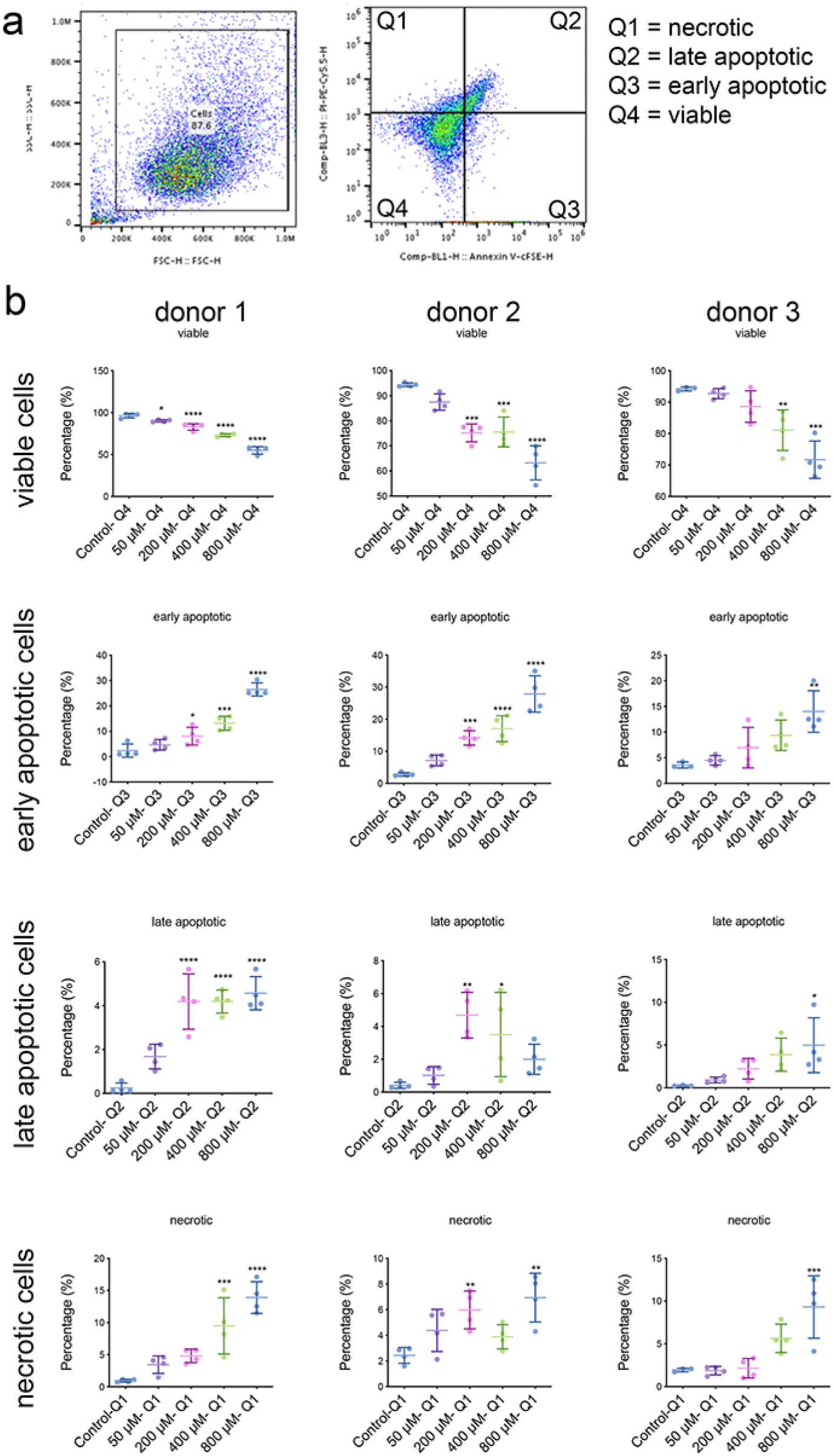
MSC viability by annexin V and propidium iodide flow cytometry with exposure to different concentrations of C12-HSL. Cells were attributed as viable, early apoptotic, late apoptotic or necrotic (graphs show mean±SD, d=3, r=4, t=1, stats by ANOVA and Tukey test where *=p<0.05, **=p<0.01, ***=p<0.001 and ****=p<0.0001).

